# Enhanced cGAS-STING-dependent interferon signaling associated with mutations in ATAD3A

**DOI:** 10.1101/2021.04.02.438201

**Authors:** Alice Lepelley, Erika Della Mina, Erika Van Nieuwenhove, Lise Waumans, Sylvie Fraitag, Gillian I Rice, Ashish Dhir, Marie-Louise Frémond, Mathieu P Rodero, Luis Seabra, Edwin Carter, Christine Bodemer, Daniela Buhas, Bert Callewaert, Pascale de Lonlay, Lien De Somer, David A Dyment, Fran Faes, Lucy Grove, Simon Holden, Marie Hully, Manju A Kurian, Hugh J McMillan, Kristin Suetens, Henna Tyynismaa, Stephanie Chhun, Timothy Wai, Carine Wouters, Brigitte Bader-Meunier, Yanick J Crow

**Affiliations:** Laboratory of Neurogenetics and Neuroinflammation, Institut Imagine, Université de Paris, Paris, France; UZ Leuven, Department of Pediatrics, Leuven, Belgium; Department of Microbiology and Immunology, Laboratory of Adaptive Immunity, KU Leuven, Leuven, Belgium; VIB-KU Leuven Center for Brain and Disease Research, Leuven, Belgium; Department of Pathology, UZ Leuven, campus Gasthuisberg, Leuven, Belgium; Service d’anatomo-pathologie, Hôpital Necker Enfants-Malades, APHP, Paris, France; Division of Evolution and Genomic Sciences, School of Biological Sciences, Faculty of Biology, Medicine and Health, University of Manchester, Manchester Academic Health Science Centre, Manchester, United Kingdom; Centre for Genomic and Experimental Medicine, MRC Institute of Genetics and Molecular Medicine, University of Edinburgh, Edinburgh, United Kingdom; Department of Dermatology and Reference Centre for Genodermatoses and Rare Skin Diseases (MAGEC), Imagine Institute, Hôpital Universitaire Necker-Enfants Malades, APHP, Université Paris-Centre, Paris, France; Medical Genetics Division, Department of Specialized Medicine, McGill University Health Centre, Montreal, Canada; Human Genetics Department, McGill University, Montreal, Canada; Center for Medical Genetics, Ghent University Hospital, Ghent, Belgium; Department of Biomolecular Medicine, Ghent University, Ghent, Belgium; Reference Center for Inherited Metabolic Diseases, Necker Hospital, APHP, INSERM, Unit 1151, INEM, Université de Paris, Filière G2M, MetabERN, Paris, France; Institut Imagine, INSERM U1163, Paris, France; Pediatric Rheumatology, UZ Leuven, Leuven, Belgium; Laboratory of Immunobiology, Rega Institute, KU Leuven, Leuven, Belgium; European Reference Network for Rare Immunodeficiency, Autoinflammatory and Autoimmune Diseases (RITA) at University Hospital Leuven, Leuven, Belgium; Department of Genetics, Children’s Hospital of Eastern Ontario, Ottawa, Ontario, Canada; Children’s Hospital of Eastern Ontario Research Institute, Ottawa, Ontario, Canada; Department of pediatric neurology, Ghent University Hospital, Ghent, Belgium; Community Paediatric Department, West Suffolk Hospital Foundation Trust, Bury St Edmunds, United Kingdom; Department of Clinical Genetics, Addenbrooke’s Hospital, Cambridge, United Kingdom; Pediatric Neurology Department, Hôpital Necker-Enfants Malades, AP-HP, Paris, France; Developmental Neurosciences, UCL GOS Institute of Child Health, London, United Kingdom; Children’s Hospital of Eastern Ontario Research Institute, University of Ottawa, Ottawa, Canada; Department of radiology, University Hospitals Leuven, Radiology, Leuven, Belgium; Department of radiology, Regional Hospital Heilig Hart Leuven, Leuven, Belgium; Stem Cells and Metabolism Research Program, Faculty of Medicine and Neuroscience Center, Helsinki Institute of Life Science, University of Helsinki, Helsinki, Finland; Paris Descartes University, Université de Paris, Sorbonne-Paris-Cité, Paris, France; Laboratory of Immunology, Hôpital Necker-Enfants Malades, Assistance Publique– Hôpitaux de Paris, Centre-Université de Paris, Paris, France; Institut Necker-Enfants Malades, Centre National de la Recherche Scientifique UMR8253, Institut National de la Santé et de la Recherche Médicale UMR1151, Team Immunoregulation and Immunopathology, Paris, France; Mitochondrial Biology Group, Institut Pasteur CNRS UMR 3691, Paris, France; Pediatric Immunology-Hematology and Rheumatology Unit, Hôpital Necker-Enfants Malades, Laboratory of Immunogenetics of Pediatric Autoimmunity, INSERM UMR 1163, APHP, Institut Imagine, Paris, France

## Abstract

Mitochondrial DNA (mtDNA) has been suggested to drive immune system activation, but the induction of interferon signaling by mtDNA has not been demonstrated in a Mendelian mitochondrial disease. We initially ascertained two patients, one with a purely neurological phenotype, and one with features suggestive of systemic sclerosis in a syndromic context, and found them both to demonstrate enhanced interferon-stimulated gene (ISG) expression in blood. We determined each to harbor a previously described de novo dominant-negative heterozygous mutation in *ATAD3A*, encoding ATPase family AAA domain-containing protein 3A (ATAD3A). We identified five further patients with mutations in *ATAD3A*, and recorded up-regulated ISG expression and interferon alpha protein in four of them. Knockdown of *ATAD3A* in THP-1 cells resulted in increased interferon signaling, mediated by cyclic GMP-AMP synthase (cGAS) and stimulator of interferon genes (STING). Enhanced interferon signaling was abrogated in THP-1 cells and patient fibroblasts depleted of mtDNA. Thus, mutations in the mitochondrial membrane protein ATAD3A define a novel type I interferonopathy.

**Summary:** Dominant-negative mutations in ATAD3A, a ubiquitously expressed mitochondrial protein, cause mitochondrial DNA-dependent up-regulation of type I interferon signaling in the context of neurological disease and autoimmunity, thereby defining a novel type I interferonopathy.

---

The type I interferonopathies are Mendelian inborn errors of immunity in which type I interferon signaling is enhanced and considered directly relevant to disease causation (Crow and Manel, 2015). Definition of this grouping has highlighted the risks to human health of the sensing of self-derived nucleic acids, leading to the induction of an inappropriate type I interferon-mediated antiviral response (Uggenti et al., 2019). Interferon has also been implicated in the pathology of a variety of complex, autoimmune disorders (Muskardin and Niewold, 2018). Mitochondria represent vestigial bacteria living in the cytosol of eukaryotic cells; and whilst they play a central role in oxidative phosphorylation, metabolism and apoptosis, there is increasing recognition of the potential of mitochondrial-derived nucleic acid to act as an agonist of the interferon signaling machinery in the context of disease (Dhir et al., 2018; Kim et al., 2019; Yu et al., 2020).

Here we report dominant-negative mutations in ATAD3A, a ubiquitously expressed mitochondrial protein, to cause mitochondrial DNA (mtDNA)-dependent up-regulation of type I interferon signaling. ATAD3A has been described to have pleiotropic functions, including in mtDNA maintenance, mitophagy dynamics and steroid synthesis (Baudier, 2018), none of which have been directly linked to type I interferon signaling. As a member of the AAA ATPase family, the protein has been suggested to form oligomers required for processive ATP hydrolysis within the oligomer (Gilquin et al, 2010), so that ATAD3A variants can act in a dominant negative manner in the presence of wildtype ATAD3A by interfering with oligomerization or ATPase activity (Cooper et al., 2017; Frazier et al., 2020). Whilst previously described patients with mutations in ATAD3A were ascertained on the basis of neurological involvement, two of the individuals that we report demonstrate features of systemic sclerosis, an autoimmune connective tissue phenotype where a disturbance of type I interferon signaling has been suggested to play an important pathogenic role (Skaug and Assassi, 2020).

## Results and Discussion

### Description of patients, their mutational status and ex vivo interferon signaling

As part of an ongoing protocol involving the agnostic screening of patients with uncharacterized phenotypes for up-regulation of type I interferon signaling, we identified two individuals demonstrating persistently enhanced interferon-stimulated gene (ISG) expression in their blood.

Patient 1 exhibits purely neurological disease (see Materials and methods for full clinical details). Now, aged seven years, she is profoundly developmentally delayed, centrally hypotonic with a peripheral dystonia and no useful motor function (Table 1). Communication is limited to eye movements and facial expression. In light of the prominent dystonia, and a raised level of neopterin in cerebrospinal fluid (CSF) suggestive of an inflammatory process, ISG expression was assessed in blood and found to be markedly elevated on each of four occasions tested between the ages of two and nine years. Trio exome sequencing identified a previously described heterozygous c.1582C>T (p.(Arg528Trp); R528W) de novo mutation in *ATAD3A* (Harel et al., 2016).

**Table 1.**
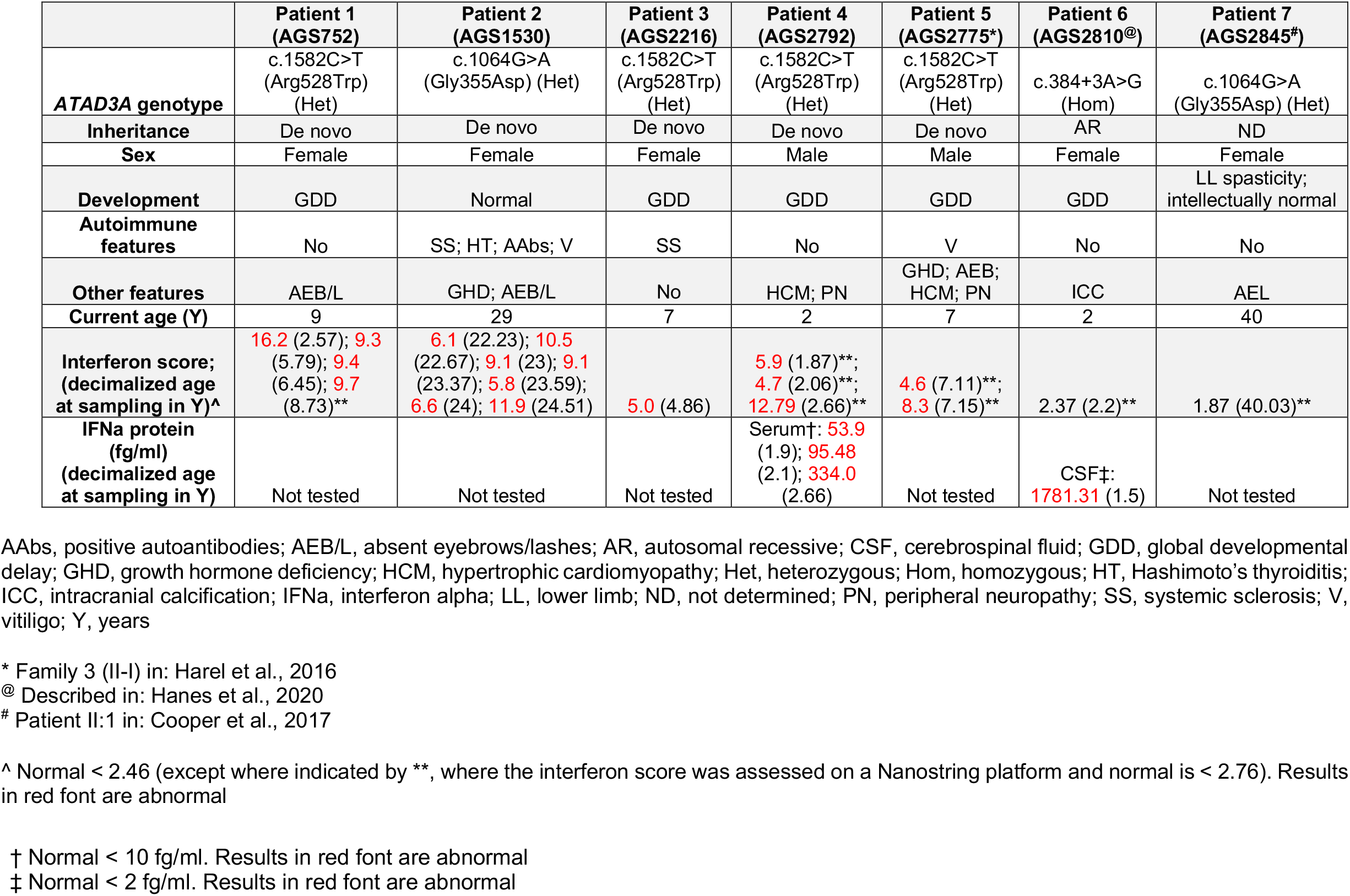
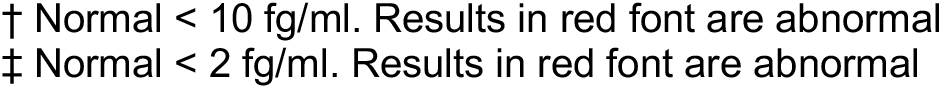
Molecular, clinical and interferon data relating to Patients 1 – 7.

|  | **Patient 1**<br>**(AGS752)** | **Patient 2**<br>**(AGS1530)** | **Patient 3**<br>**(AGS2216)** | **Patient 4**<br>**(AGS2792)** | **Patient 5**<br>**(AGS2775\*)** | **Patient 6**<br>**(AGS2810@)** | **Patient 7**<br>**(AGS2845#)** |
| --- | --- | --- | --- | --- | --- | --- | --- |
| **ATAD3A genotype** | c.1582C>T<br>(Arg528Trp)<br>(Het) | c.1064G>A<br>(Gly355Asp) (Het) | c.1582C>T<br>(Arg528Trp)<br>(Het) | c.1582C>T<br>(Arg528Trp)<br>(Het) | c.1582C>T<br>(Arg528Trp)<br>(Het) | c.384+3A>G<br>(Hom) | c.1064G>A<br>(Gly355Asp) (Het) |
| **Inheritance** | De novo | De novo | De novo | De novo | De novo | AR | ND |
| **Sex** | Female | Female | Female | Male | Male | Female | Female |
| **Development** | GDD | Normal | GDD | GDD | GDD | GDD | LL spasticity;<br>intellectually normal |
| **Autoimmune**<br>**features** | No | SS; HT; AAbs; V | SS | No | V | No | No |
| **Other features** | AEB/L | GHD; AEB/L | No | HCM; PN | GHD; AEB;<br>HCM; PN | ICC | AEL |
| **Current age (Y)** | 9 | 29 | 7 | 2 | 7 | 2 | 40 |
| **Interferon score;**<br>**(decimalized age**<br>**at sampling in Y)^** | 16.2 (2.57); 9.3<br>(5.79); 9.4<br>(6.45); 9.7<br>(8.73)** | 6.1 (22.23); 10.5<br>(22.67); 9.1 (23); 9.1<br>(23.37); 5.8 (23.59);<br>6.6 (24); 11.9 (24.51) | 5.0 (4.86) | 5.9 (1.87)**;<br>4.7 (2.06)**;<br>12.79 (2.66)** | 4.6 (7.11)**;<br>8.3 (7.15)** | 2.37 (2.2)** | 1.87 (40.03)** |
| **IFNa protein**<br>**(fg/ml)**<br>**(decimalized age**<br>**at sampling in Y)** | Not tested | Not tested | Not tested | Serum†: 53.9<br>(1.9); 95.48<br>(2.1); 334.0<br>(2.66) | Not tested | CSF‡:<br>1781.31 (1.5) | Not tested |

AAbs, positive autoantibodies; AEB/L, absent eyebrows/lashes; AR, autosomal recessive; CSF, cerebrospinal fluid; GDD, global developmental delay; GHD, growth hormone deficiency; HCM, hypertrophic cardiomyopathy; Het, heterozygous; Hom, homozygous; HT, Hashimoto's thyroiditis; ICC, intracranial calcification; IFNa, interferon alpha; LL, lower limb; ND, not determined; PN, peripheral neuropathy; SS, systemic sclerosis; V, vitiligo; Y, years
\* Family 3 (II-I) in: Harel et al., 2016
@ Described in: Hanes et al., 2020

Patient 2 attended university with no intellectual deficits, but demonstrates a spastic diplegic gait which appears stable since early infancy. She was diagnosed with antibody-positive Hasimoto’s thyroiditis, vitiligo and growth hormone deficiency in childhood (Table 1). Beginning at 21 years of age she developed sclerodermatous involvement of the hands, face and ventral surface of the forearms, which progressed rapidly over a few months (Rodnan skin score: 33/51), fulfilling criteria for a diagnosis of systemic sclerosis (van den Hoogen et al., 2013) (Fig. S1 A). Skin biopsy of affected tissue showed classical features of systemic sclerosis (Fig. S1 B). Immunoglobulin levels, antinuclear and anti-double stranded (ds) DNA antibody titers were persistently elevated, but systemic sclerosis-related antibodies were not detected (Table S1). Continued clinical progression despite oral corticosteroids led to the introduction of rapamycin, which was associated with a partial regression of the cutaneous fibrosis and a decrease of the Rodnan skin score to 10/51 over two years. ISG expression in blood was markedly elevated on each of seven occasions tested between the ages of 22 and 24 years. Trio exome sequencing identified a previously described heterozygous c.1064G>A (p.(Gly355Asp); G355D) de novo mutation in *ATAD3A* (Cooper et al., 2017).

Given the above findings, we went on to ascertain other patients known to harbor mutations in *ATAD3A*. In this way, we recruited a further five individuals (Table 1 and Materials and methods).

Patient 3 demonstrated early onset severe motor delay with impaired cognition but good social contact. From age two years she developed progressive hardening of the skin. At four years of age she exhibited peri-oral sclerosis, tight sclerodermatous skin of the upper and lower limbs, and severe flexion contractures at the knees, ankles, elbows, wrists and fingers (Rodnan skin score: 32/51) (Fig. S1 A). Skin biopsy revealed features typical of systemic sclerosis (Fig. S1 B). MRI imaging of the upper limbs confirmed cutaneous thickening (Fig. S1 C). Treatment with rapamycin and low dose corticosteroids was associated with a good clinical response. She was shown to carry the same p.(Arg528Trp) substitution recorded in Patient 1. She demonstrated enhanced ISG expression in blood on the single occasion tested at age four years.

Patients 4 and 5, both males, exhibit severe global developmental delay, hypertrophic cardiomyopathy and a peripheral neuropathy. Patient 5, like Patient 2, has a history of vitiligo and growth hormone deficiency. Each of Patient 4 and Patient 5 carries the same p.(Arg528Trp) substitution in ATAD3A as seen Patients 1 and 3, Patient 5 having been previously included in the first published series of cases with mutations in *ATAD3A* (Family 3 II-I in (Harel et al., 2016)). Both demonstrated enhanced expression of ISGs in blood (three and two times respectively). Additionally, when assessed at the same time as ISG expression, elevated levels of interferon alpha protein were recorded three times in the serum of Patient 4.

As recently reported (Hanes et al., 2020), Patient 6, a female, has severe neurological disease due to a homozygous pathogenic splice-site transition (c.528+3A>G) in *ATAD3A*. CSF interferon alpha protein was highly elevated at one year of age, whilst blood ISG expression was normal on the one occasion tested at age two years. Bilateral calcification of the basal ganglia was present on cranial CT at this time.

Patient 7 is of completely normal intellect into mid-adulthood. As previously identified by Cooper et al. (Cooper et al., 2017), she carries the same p.(Gly355Asp) mutation in ATAD3A as seen in Patient 2. She was noted to demonstrate bilateral lower limb spasticity in early childhood. At age 40 years she exhibits severe lower limb spasticity and weakness. ISG expression was normal when assessed at this age. Although not tested here, her son, carrying the same mutation as his mother, has a severe dyskinetic motor disorder and profound intellectual delay.

### Summary of molecular data

Six of seven patients ascertained in this study are heterozygous for one of two missense mutations (p.(Gly355Asp); p.(Arg528Trp)) in ATAD3A (Fig. 1 A), neither of which is recorded on publicly available databases, being absent from >250,000 control alleles on gnomAD. The substituted amino acids are highly evolutionarily conserved (Fig. 1 B), and the amino acid substitutions are predicted as damaging by in silico algorithms (SIFT: Damaging; Polyphen2: Probably damaging). The glycine to aspartate substitution at amino acid 355 falls within the highly conserved Walker A motif, responsible for ATP binding in the AAA+-ATPase module of ATAD3A (Fig. 1C) (Baudier, 2018). Importantly, recombinant p.(Gly355Asp) mutant ATAD3A protein was shown to be associated with a markedly reduced ATPase activity, and demonstrated a strong dominant-negative effect on the ATPase activity of wild-type protein (Cooper et al., 2017). Similarly, the Drosophila equivalent of human p.(Arg528Trp) has also been documented to behave as a dominant negative allele (Harel et al., 2016). Meanwhile, the homozygous c.528+3A>G transition seen in Patient 6 results in a significant reduction of ATAD3A protein in patient fibroblasts (Hanes et al., 2020).

**Figure 1.**
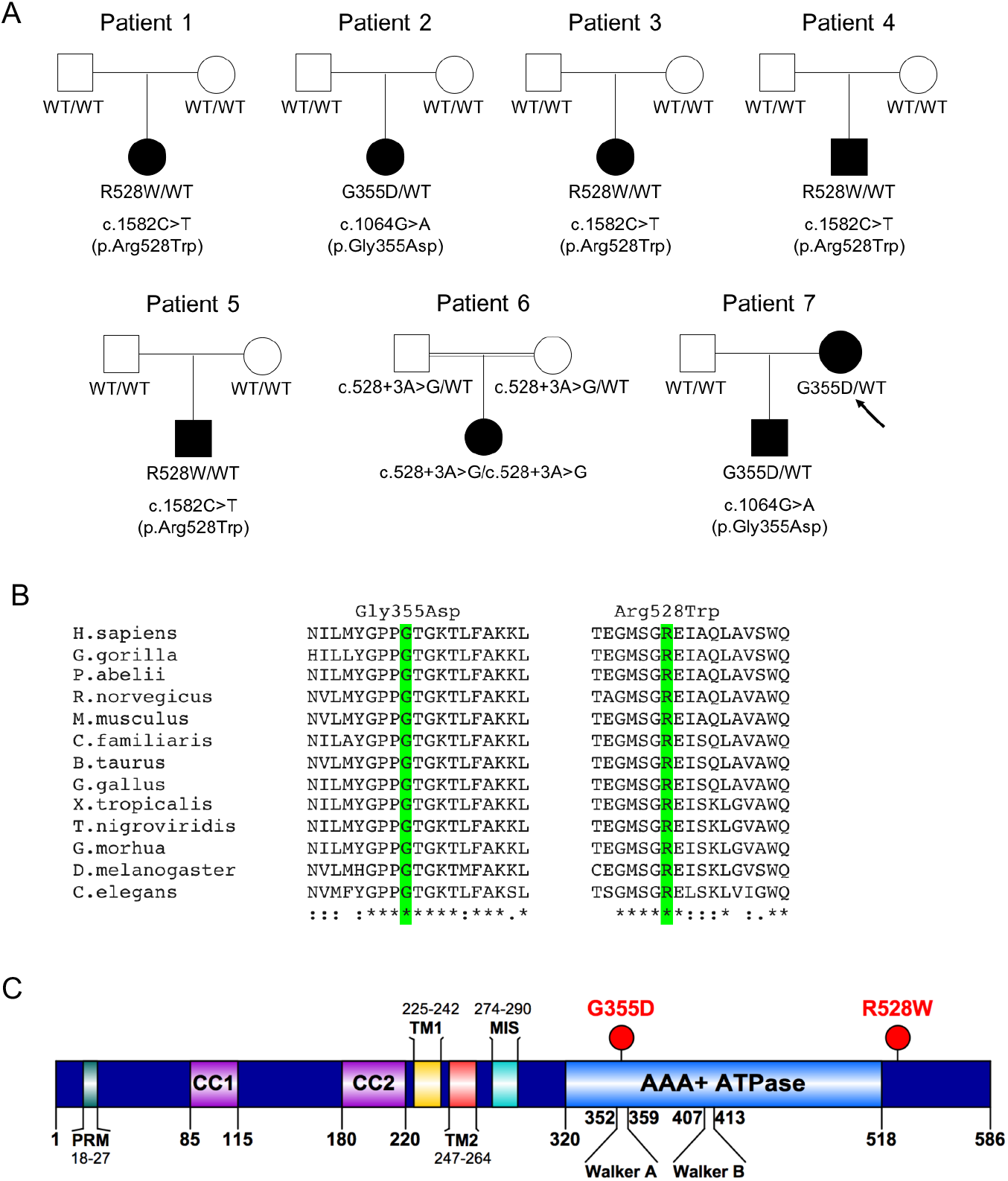
*ATAD3A* genetic data. (A) Family structures relating to Patients 1 -7. Solid symbols indicate the affected patients, open symbols unaffected parents, squares males, circles females, double-line consanguinity. Genotype is indicated below each individual (WT: wild-type). Genomic mutation (NM_001170535) and amino acid changes are detailed below each family. Arrow indicates Patient 7. (B) Evolutionary conservation of the ATAD3A amino acid substitutions p.(Gly355Asp) (G355D) and p.(Arg528Trp) (R528W) (highlighted in green). (C) Domains of the ATAD3A protein, and the position of the p.(Gly355Asp) and p.(Arg528Trp) substitutions. G355 is located in the ATP binding (Walker A) subdomain of the AAA+ ATPase domain. R528 is located in the C-terminus of ATAD3A after the AAA+ ATPase domain. PRM denotes a proline-rich motif, CC coiled-coil domain, TM transmembrane domain, MIS mitochondrial import signal (adapted from Baudier, 2018). The homozygous c.528+3A>G spice site variant identified in Patient 6, leading to a loss of ATAD3A protein expression, is not depicted.

### Summary of ex vivo interferon signaling

The combination of neurological and immunological features is characteristic of diseases within the type I interferonopathy grouping, and patients with type I interferonopathies demonstrate high levels of ISG messenger RNA in whole blood. Using a previously validated assay (Rice et al., 2013), we recorded an increased expression of ISGs in peripheral blood in five of the seven patients assessed. Importantly, positive serial data were available in four patients, including two patients sampled seven and four times over a period of more than 2 and six years respectively (Table 1; Fig. 2 A). Although in two probands ISG expression was not increased on the single occasion tested, we observed a markedly increased level of interferon alpha protein in the CSF of one of these individuals.

**Figure 2.**
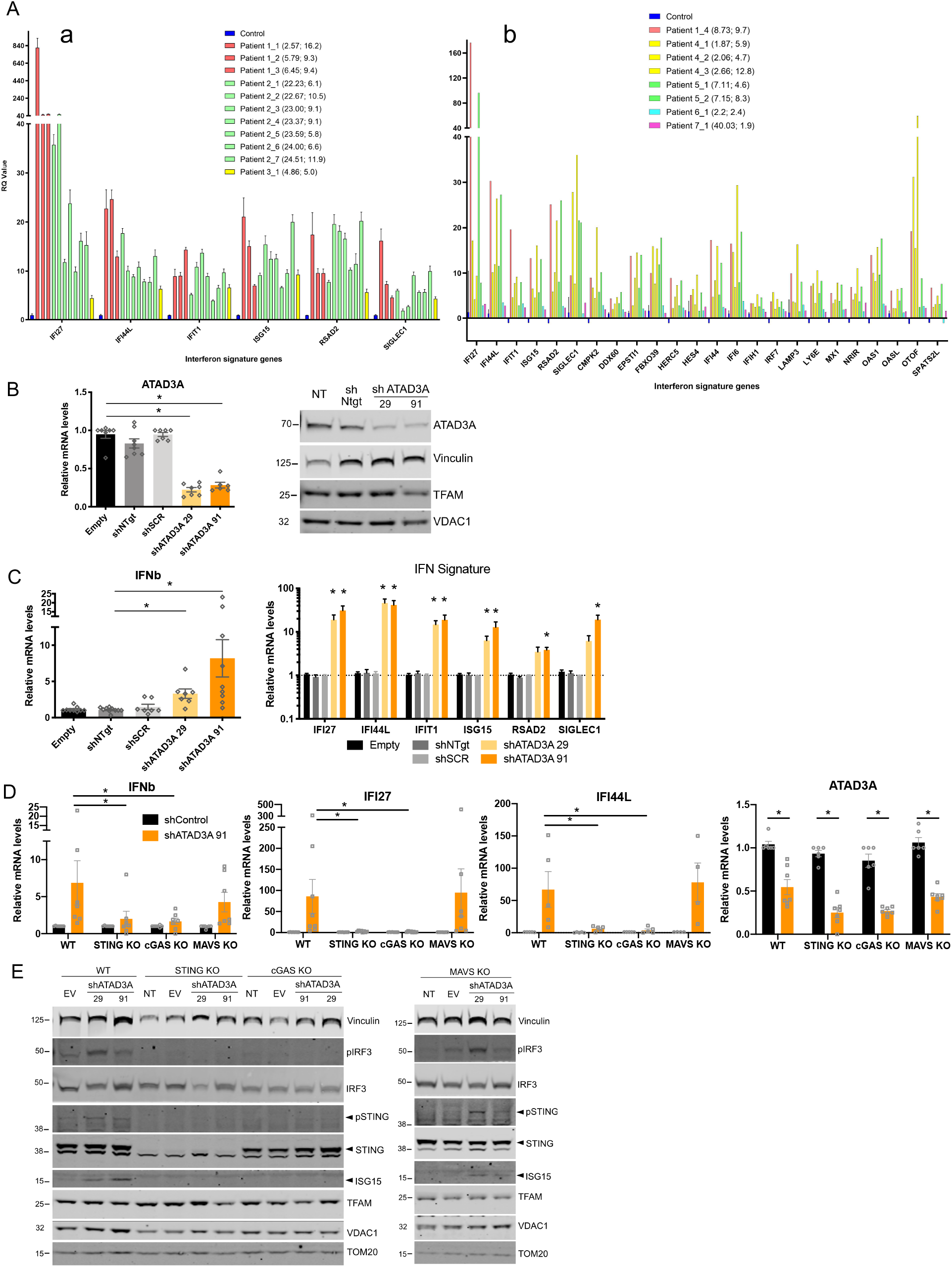
ATAD3A dysfunction leads to up-regulated interferon signaling in patient blood and in THP-1 cells through the cGAS-STING pathway. (A) Enhanced interferon-stimulated gene (ISG) messenger RNA expression in peripheral whole blood recorded in five of seven patients with *ATAD3A* mutations. Subpanel a shows the expression of six ISGs measured using TaqMan probes, and subpanel b a larger panel of 24 genes assessed using a Nanostring platform. Colored bars indicate different patients. The first number in the brackets is the decimalized age at the time of sampling, and the second the interferon score (calculated according to the median fold change in relative quantification of the ISGs compared to 29 (a) or 27 (b) controls (blue bars) (normal < 2.466 (a) or < 2.75 (B)). Knockdown of *ATAD3A* gene expression (qPCR) (left) and of ATAD3A protein (western blot) (right) in THP-1 cells using two short-hairpin (sh) RNAs (shATAD3A 29 and shATAD3A 91) compared to controls (transduction with a vector with no shRNA (empty/EV), or two different unspecific scrambled shRNAs (shNTgt and shSCR)), or non-transduced (NT), and analyzed 4 days later. For qPCR, mean values of six to seven independent experiments are shown and are expressed as the ratio of the mRNA levels, normalized to housekeeping gene *HPRT* mRNA, in indicated conditions to that in the empty condition. * indicates statistical significance using a Kruskal Wallis test with Dunn’s post-hoc. Western blot images are representative of 4 experiments. VDAC1 and TFAM are mitochondrial proteins, Vinculin is a loading control. (C) Knockdown of *ATAD3A* leads to increased expression of the interferon beta (*IFNb*) gene, and of six representative ISGs (*IFI27, IFI44L, IFIT1, ISG15, RSAD2, SIGLEC1*), compared to control vectors (EV, shNTgt and shSCR), in cells treated as in (B). Mean values of seven to ten independent experiments are shown and are expressed as the ratio of the mRNA levels, normalized to housekeeping gene *HPRT* mRNA, in indicated conditions to that in the empty condition. For *IFNb*, * indicates significance in Kruskal Wallis test with Dunn’s post-hoc. For ISGs, * indicates statistical significance in 2-way ANOVA with Dunnett’s multiple comparison test to shSCR. (D) Knockdown of *ATAD3A* with shATAD3A 91 in wild-type (WT) THP-1 cells and in cells null for either *cGAS, STING* or *MAVS*. The increased expression of interferon beta (*IFNb*) and two representative ISGs (*IFI27, IFI44L*) seen in WT THP-1 cells is lost in cells that are null (KO) for either *cGAS* or *STING*, whilst THP-1 cells null for *MAVS* behave similar to WT cells. Mean values of seven to eight independent experiments are shown and are expressed as the ratio of the mRNA levels, normalized to housekeeping gene *HPRT* mRNA, in indicated conditions to that with control shRNA (‘shControl’: shNTgt or shSCR). * indicates statistical significance using 2-way ANOVA with Dunnett’s multiple comparison test. (E) Western blot analysis of *ATAD3A* knockdown compared to untreated (NT) cells, or cells treated with an empty vector (EV), showing an increase in the expression of phosphorylated IRF3, phosphorylated STING and ISG15 (bottom band) protein in WT and MAVS KO cells, but not in STING KO or cGAS KO cells. One experiment representative of three is shown. TOM20, VDAC1 and TFAM are mitochondrial proteins, Vinculin is a loading control.

### Type I interferon signaling in THP-1 cells upon ATAD3A knockdown

Given previous data showing that the two recurrent ATAD3A missense substitutions identified in our patients act as dominant-negative alleles (Cooper et al., 2017; Harel et al., 2016), we performed ATAD3A silencing by lentiviral transduction of THP-1 cells with two different short hairpin RNAs (shRNAs) targeting *ATAD3A*. Four days after transduction, knockdown of *ATAD3A* messenger RNA resulted in a decrease in ATAD3A protein (Fig. 2 B), and an increase in interferon beta and representative ISG expression (*IFI27, IFI44L, IFIT1, ISG15, RSAD2, SIGLEC1*) (Fig. 2 C). We also observed increased IRF3 phosphorylation, a key event upstream of interferon beta induction, and ISG15 protein expression (Fig. 2 E). These data were confirmed by CRISPR editing, leading to down-regulation of *ATAD3A* in THP-1 cell pools using two different guide RNAs, where we also recorded increased phosphorylated IRF3 upon loss of *ATAD3A* (Fig. S2, A-D).

### Definition of the pathway mediating type I interferon signaling upon ATAD3A knockdown

In mammalian cells, immune responses to viral infection involve host-encoded nucleic acid-binding pattern recognition receptors, including the cytosolic sensors of dsRNA (melanoma differentiation-associated protein 5 (MDA5) and retinoic acid-inducible gene I (RIG-I), both signaling through mitochondrial-antiviral signaling protein (MAVS), and DNA, most particularly cGAS (signaling through STING). We noticed increased phosphorylation of STING, indicative of STING activation, upon *ATAD3A* knockdown in wild-type (WT) THP-1 cells (Fig. 2 E). In order to explore the pathway mediating the observed interferon induction, we compared the results of knockdown of *ATAD3A* in WT THP-1 cells, and in characterized THP-1 cells null for *cGAS, STING* or *MAVS*. Increased STING phosphorylation and enhanced interferon signaling, evidenced by interferon beta and ISG up-regulation, IRF3 phosphorylation and ISG15 protein induction, after *ATAD3A* knockdown by shRNA and CRISPR editing, was preserved in THP-1 cells null for *MAVS* as in wild-type cells, but such enhanced signaling was abrogated in the absence of either *cGAS* or *STING* (Fig. 2 D and E and Fig. S2 E).

### Interferon signaling is dependent on the presence of mtDNA upon ATAD3A knockdown

Considering the role of ATAD3A in mtDNA maintenance, and its interaction with the mtDNA nucleoid organizer TFAM (mitochondrial transcription factor A) (He et al., 2012; Zhao et al., 2019), we suspected that cytosolic re-localization of mtDNA, and subsequent sensing by cGAS, might be driving interferon stimulation (West et al., 2015). To explore this further, we analyzed the effect of the knockdown of *ATAD3A* in THP-1 cells transiently depleted of mtDNA (Fig. 3 A). Compared to untreated THP-1 cells, knockdown of *ATAD3A* in mtDNA-depleted THP-1 cells induced lower levels of expression of interferon beta and ISGs (Fig. 3 B). The induction of IRF3 phosphorylation and ISG15 protein by *ATAD3A* knockdown was also abrogated by mtDNA depletion (Fig. 3 C). As expected, knockdown of *TFAM* led to enhanced interferon signaling (considered due to mtDNA cytosolic leakage: (West et al., 2015)), which was again lost in mtDNA-depleted THP-1 cells (Fig. 3, B and C).

**Figure 3.**
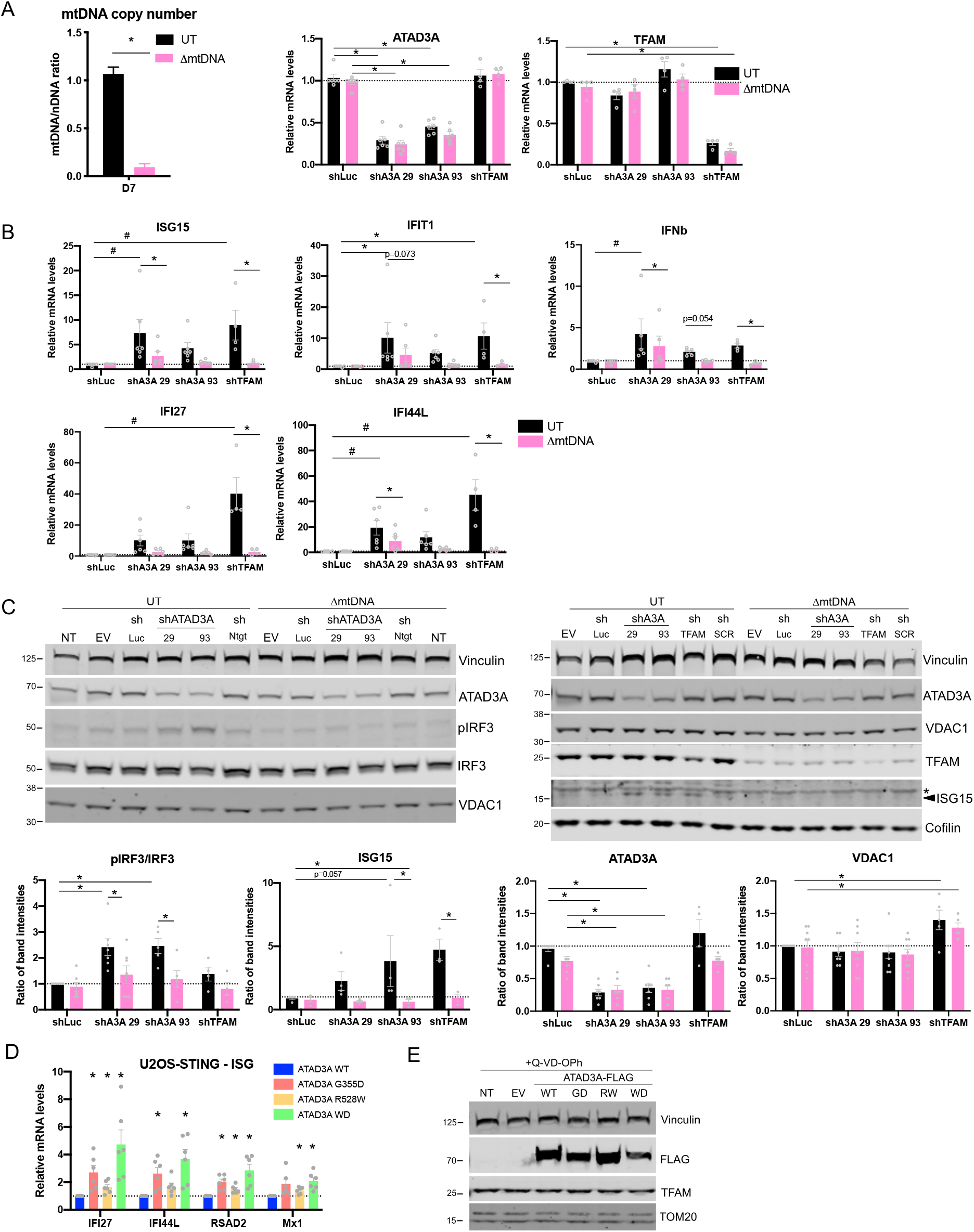
Interferon signaling upon ATAD3A silencing in THP-1 cells is dependent on the presence of mtDNA. (A) Depletion of mtDNA in THP-1 cells. Graph shows qPCR analysis of mtDNA copy number per cell, expressed as the ratio of the DNA quantity of the mitochondrial gene (*MT-COXII*) over the nuclear gene (*GAPDH*), in total DNA isolated from untreated (UT) THP-1 cells, or cells treated for 7 days with low dose ethidium bromide to deplete mtDNA (ΔmtDNA). The average of four independent experiments is shown. * Indicates statistical significance using a Mann-Whitney test. (B) Knockdown of *ATAD3A* assessed by qPCR using either of two shRNAs (shATAD3A 29, shATAD3A 93), or of *TFAM* using shTFAM, in mtDNA-depleted THP-1 cells compared to control shRNA (shLuc). Upon knockdown of either *ATAD3A* or *TFAM*, the increase in the expression of interferon beta (*IFNb*) and of four representative ISGs (*ISG15, IFIT1, IFI27, IFI44L*) seen in untreated (UT) THP-1 cells was lost when THP-1 cells depleted of mtDNA were used. Mean values of five to six independent experiments are shown, and are expressed as the ratio of the mRNA levels, normalized to housekeeping gene *HPRT* mRNA, in indicated conditions to that in the control shLuc condition. For *ATAD3A* and *TFAM* expression, * indicates statistical significance in 2-way ANOVA with Dunnett’s multiple comparison test compared to respective shLuc. For *IFNb* and ISGs, # indicates statistical significance in 2-way ANOVA with Dunnett’s multiple comparison test compared to respective shLuc, and * indicates statistical significance in 2-way ANOVA with Sidak’s multiple comparison test comparing ΔmtDNA and UT conditions for each shRNA. (C) Western blot analysis of *ATAD3A* and *TFAM* knockdowns described in (B), compared to untreated (NT) cells, or cells treated with an empty vector (EV) or a control shRNA (shNTgt/shLuc), showing an increase in the expression of phosphorylated IRF3 (left) and ISG15 (right, bottom band; * indicates non-specific band) protein; abrogated by mtDNA depletion. Two representative experiments are shown. Quantification of band intensities is expressed as the ratio of phosphorylated IRF3/total IRF3 or the ratio of ISG15, ATAD3A and VDAC1 (mitochondrial protein control) protein levels normalized to Vinculin, averaged from four to seven independent experiments. * indicates statistical significance in 2-way ANOVA with Sidak’s multiple comparison test comparing relevant shRNA to control shLuc in each cell line, and comparing UT and mtDNA-depleted cells for each shRNA. (D) mRNA expression analysis assessed by qPCR of ISGs *IFI27, IFI44L, RSAD2* and *Mx1* in U2OS cells stably expressing STING (U2OS-STING), transfected with ATAD3A WT or mutant G355D, R528W and WD plasmids, treated with 10 μM QVD-OPh after 24 hours and harvested 24 hours later. Expression levels are expressed as the fold of levels in WT transfected cells. Means ± SEM of six independent experiments, statistically analyzed using 2-way ANOVA and Dunnett’s multiple comparisons test. (E) Western blot analysis of FLAG, Vinculin, TFAM and TOM20 in whole-cell lysates of U2OS-STING treated and harvested as in (D). Representative results from four independent experiments.

### Type I interferon signaling upon mutant ATAD3A overexpression

In trying to assess the functional effect of the p.(Gly355Asp) and p.(Arg528Trp) substitutions in the context of interferon signaling, we observed that overexpression of ATAD3A (wild-type or mutant) significantly compromised cell survival in THP-1 cells, U2OS cells and primary healthy fibroblasts (data not shown), consistent with previous reports of defects induced by overexpression of wild-type AAA ATPases (Evans et al., 2005; Harel et al., 2016). To circumvent this problem, we expressed these mutations in U2OS cells stably expressing STING and endogenous cGAS (Chen et al., 2020), and treated the cells with the caspase inhibitor Q-VD-OPh so as to block caspase-mediated apoptosis (Rongvaux et al., 2014). In this way we were able to avoid cell death secondary to ATAD3A overexpression, and observed increased ISG expression with both mutations (Fig. 3, D and E). Importantly, a similar effect was seen with a dominant negative Walker A dead (WD) K358A mutation, suggesting a dependency on the ATP-binding function of ATAD3A, and confirming a dominant negative action of ATAD3A mutations on interferon signaling.

### ATAD3A patient-derived cells show enhanced interferon signaling dependent on DNA sensing and the presence of mtDNA

Our knockdown approaches in THP-1 suggested that loss of ATAD3A function leads to interferon signaling due to DNA sensing of mtDNA. In primary human fibroblasts, ATAD3A knockdown also led to ISG induction (Fig. 4, A and B), dependent on cGAS (Fig. 4 C). Interestingly, this was independent of TFAM expression (Fig. 4, A and B). To verify these findings in the context of the disease-causing dominant-negative mutations, we assessed interferon signaling in patient fibroblasts and peripheral blood mononuclear cells (PBMCs). ATAD3A and TFAM were expressed at levels similar in patients and controls (Fig. 4 E, Fig. S2, G and H and Fig. S3 A and B). We observed interferon beta and ISG upregulation at the mRNA and protein levels in patient-derived fibroblasts and PBMCs, albeit to different extents between patients (Fig. 4 D and E and Fig. S2 F). To simplify our readout, we derived an interferon score from the median fold change in expression of six to seven ISGs (details in figure legends). As with ATAD3A knockdown cells, interferon signaling involved cGAS DNA sensing but not MAVS-dependent RNA sensing (Fig. 4 F and Fig. S3 D). Total mtDNA levels were not increased in patient fibroblasts compared to controls (Fig. S3 C), ruling out a variation in mtDNA levels as a cause of interferon induction. As with ATAD3A knockdown in THP-1 cells, interferon signaling was abrogated after mtDNA depletion with 2′,3′-dideoxycytidine (ddC) treatment (Fig. 4, G and H and Fig. S3 E).

**Figure 4.**
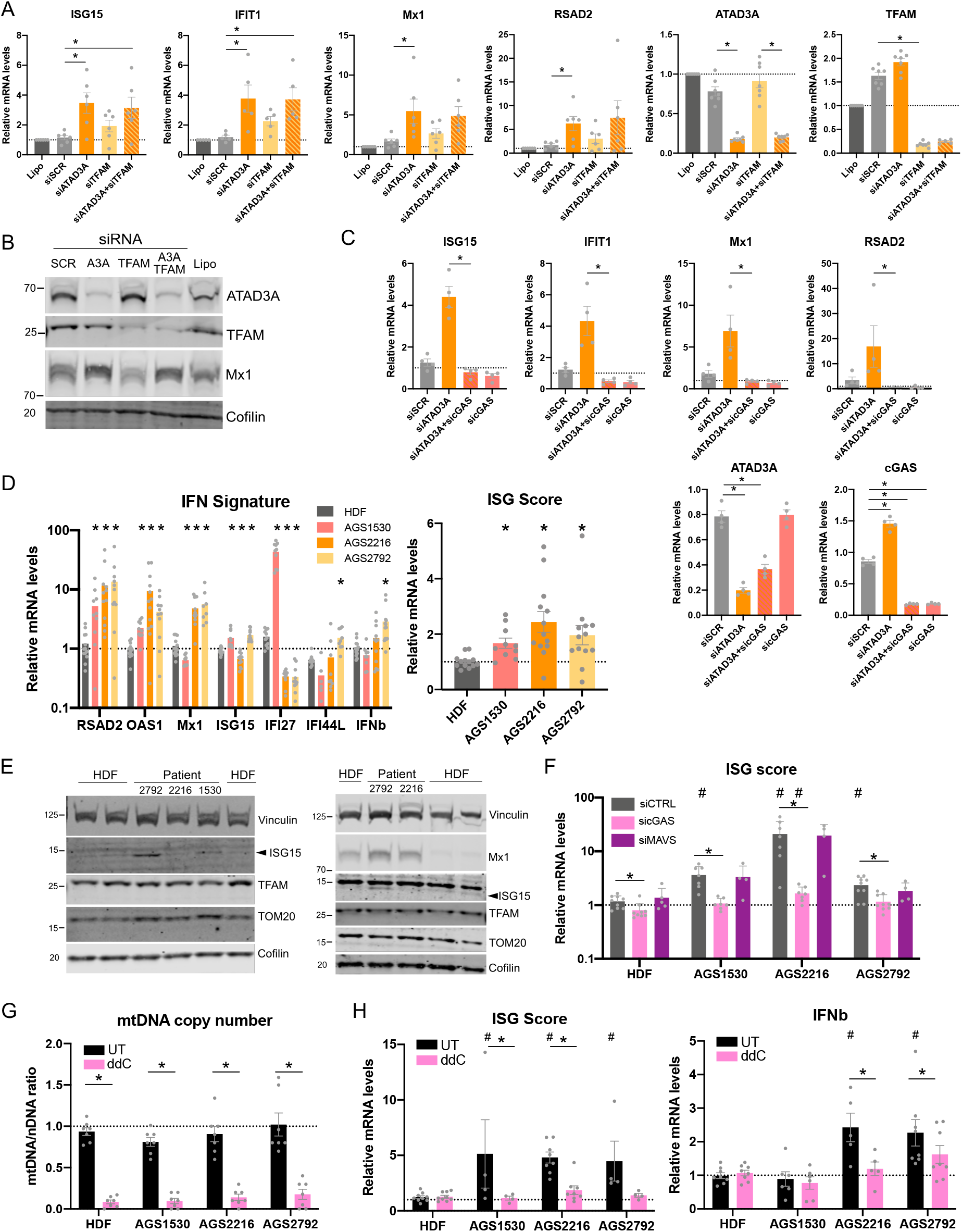
Interferon signaling in fibroblasts from ATAD3A patients is cGAS-and mtDNA-dependent. (A) ISG, *ATAD3A* and *TFAM* mRNA expression in control primary fibroblasts after downregulation of ATAD3A (siATAD3A), TFAM (siTFAM) or both (siATAD3A+siTFAM) by siRNA or treatment with control siRNA (siSCR) or transfection reagent alone (Lipo) for 72 hours, expressed as the ratio of the mRNA levels, normalized to housekeeping gene *HPRT* mRNA, in indicated conditions to that in siSCR. Mean values and data points of six to seven independent experiments are shown. * indicates statistical significance in Friedman test with Dunn’s multiple comparison test to siSCR condition. (B) Protein expression in primary fibroblasts treated as in (A) of a representative ISG Mx1, ATAD3A and TFAM assessed by western blot. Cofilin is a loading control. (C) ISGs, *ATAD3A* and *cGAS* mRNA expression in control primary fibroblasts after downregulation of ATAD3A (siATAD3A), cGAS (sicGAS) or both (siATAD3A+sicGAS) by siRNA or treatment with control siRNA (siSCR) or transfection reagent alone (Lipo) for 72 hours, expressed as in (A). Mean values and data points of six to seven independent experiments are shown, three data sets are also displayed in (A). * indicates statistical significance in Mann-Whitney test between siATAD3A and siATAD3A+sicGAS conditions for ISGs. * indicates statistical significance in one-way ANOVA for *ATAD3A* and *cGAS*. (D) Left, ISG and interferon beta *(IFNb)* mRNA expression in primary fibroblasts from control human dermal fibroblasts (HDF, average of 3) or from ATAD3A patients 2, 3 and 4 (respectively: AGS1530, AGS2216 and AGS2792), expressed as the ratio of the mRNA levels, normalized to housekeeping gene *HPRT* mRNA, in indicated conditions to that in one control HDF. Mean values and data points of ten to thirteen independent experiments are shown. * indicates statistical significance in 2-way ANOVA with Dunnett’s multiple comparison test. Right, ISG mRNA expression, represented as an ISG score (i.e. the median fold change of mRNA levels of six ISGs (*RSAD2, OAS1, Mx1, IFI27, ISG15, IFI44L*), in control and ATAD3A patient-derived primary fibroblasts shown on the left. * indicates statistical significance in Kruskal-Wallis test with Dunn’s multiple comparison (E) Protein expression in patient-derived primary fibroblasts of two representative ISGs (ISG15 and Mx1) assessed by western blot. TFAM and TOM20 are mitochondrial proteins, Cofilin and Vinculin loading controls. Arrowhead indicates relevant band for ISG15. (F) ISG mRNA expression, represented as an ISG score (i.e. the median fold change of mRNA levels of seven ISGs (*RSAD2, OAS1, Mx1, IFI27, ISG15, IRF7, IFI44L*) expressed as in (D)), in control and ATAD3A patient-derived primary fibroblasts, after downregulation of cGAS (sicGAS) or MAVS (siMAVS) by siRNA or treatment with control siRNA (siCTRL). Mean values and data points of six to nine independent experiments are shown. * indicates statistical significance in 2-way ANOVA with Dunnett’s multiple comparison tests between siRNA conditions for each cell line. # indicates statistical significance in 2-way ANOVA with Dunnett’s multiple comparison test between control and patient-derived fibroblasts for each siRNA condition. (G) Depletion of mtDNA in patient-derived primary fibroblasts. Graph shows qPCR analysis of mtDNA copy number per cell, expressed as the ratio of the DNA quantity of the mitochondrial gene (*MT-COXII*) over the nuclear gene (*GAPDH*), in total DNA isolated from untreated (UT) fibroblasts, or cells treated for 7 to 14 days with 100 μM 2′,3′ dideoxycytidine (ddC) to deplete mtDNA. The mean of seven independent experiments is shown. * indicates statistical significance in 2-way ANOVA with Sidak’s multiple comparison test comparing ddC and UT conditions for each fibroblast line. (H) ISG mRNA expression, represented as an ISG score (i.e. the median fold change of mRNA levels of seven ISGs (*RSAD2, OAS1, Mx1, IFI27, ISG15, IRF7, IFI44L*) expressed as in (D)), and interferon beta (*IFNb*) mRNA expression in control and ATAD3A patient-derived primary fibroblasts, untreated (UT) or mtDNA-depleted by ddC as in (G). Mean values and data points of three to eight independent experiments are shown. * indicates statistical significance in 2-way ANOVA with Sidak’s multiple comparison test between UT and ddC conditions. # indicates statistical significance in 2-way ANOVA with Dunnett’s multiple comparison test between control and patient-derived fibroblasts for each treatment.

### ATAD3A dysfunction leads to cytosolic release of mtDNA

The above data were consistent with interferon signaling due to the cytosolic release of mtDNA sensed by cGAS upon ATAD3A dysfunction. This phenomenon has been reported to depend on mitochondrial outer membrane opening via BAX/BAK1 macropores, VDAC1 channels or a contribution of the mitochondrial permeability transition pore mPTP (Kim et al., 2019; McArthur et al., 2018; Riley et al., 2018; Yu et al., 2020). Although we did not detect any effect of downregulating BAX and BAK1 or inhibiting their oligomerization with BAX inhibitor peptide V5 (Fig. S3, G-I), we observed that VDAC1 inhibition with 4,4′-diisothiocyanostilbene-2,2′-disulfonic acid (DIDS) led to a partial rescue of interferon beta and ISG upregulation in ATAD3A patient fibroblasts (Fig. 5 A and Fig. S3 F). Notably, whilst we did not observe any abnormality of mitochondrial network morphology on confocal microscopy in patient fibroblasts (Fig. 5, B and C), downregulation of ATAD3A in fibroblasts led to the accumulation of cytosolic DNA foci observed by confocal microscopy after staining dsDNA and mitochondria (Fig. 5 D), reminiscent of mtDNA released into the cytosol (McArthur et al., 2018; Yu et al., 2020). Rapamycin is an anti-inflammatory drug which showed a beneficial effect on the features of systemic sclerosis in Patients 2 and 3. As an inhibitor of mTOR, rapamycin is also an inducer of general autophagy, including mitophagy, the removal of damaged mitochondria (Li et al., 2014). Of interest then, we observed a marked decrease of ISG and interferon beta expression in patient fibroblasts following rapamycin treatment (Fig. 5 A and Fig. S3 F).

**Figure 5.**
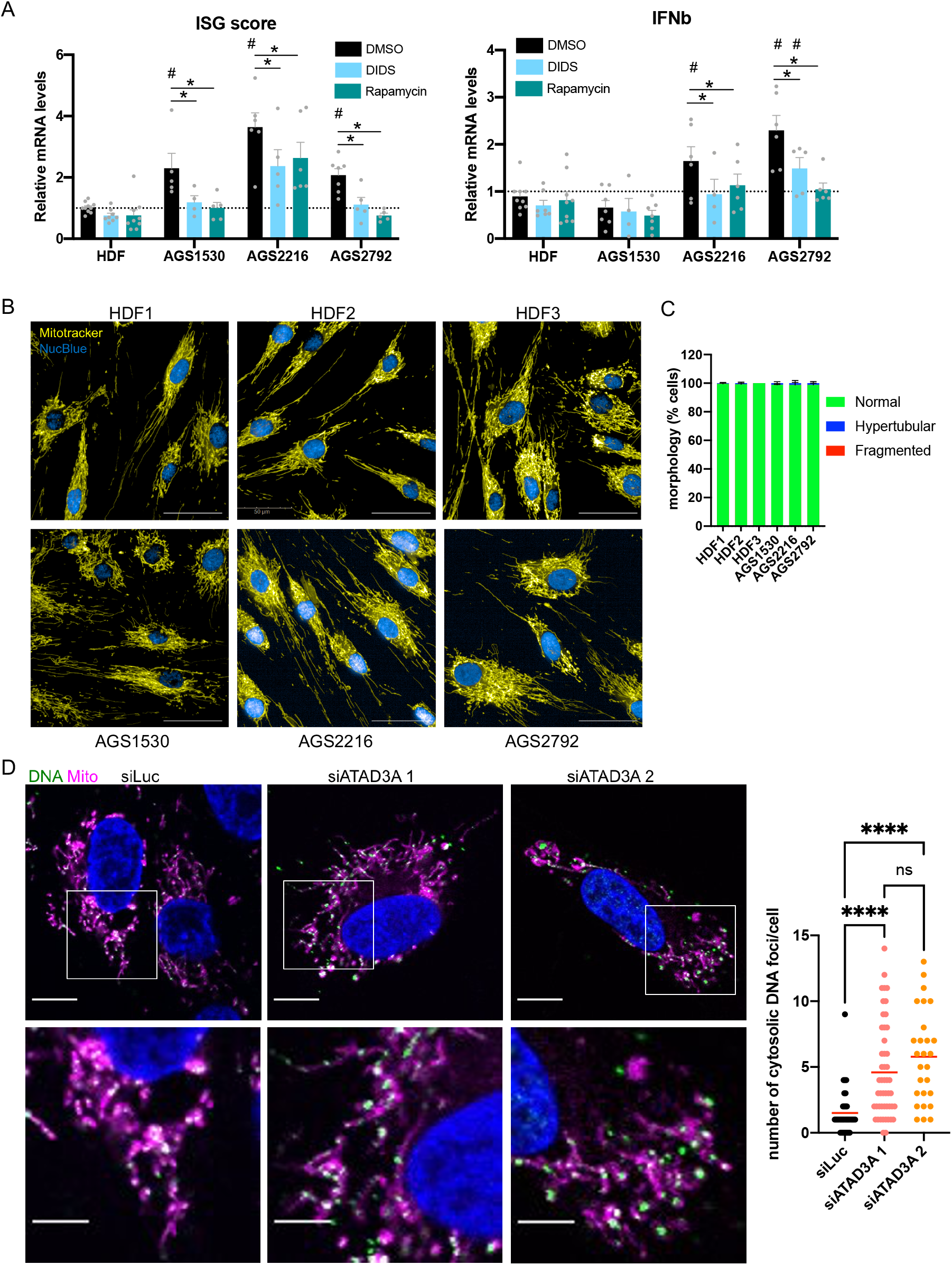
ATAD3A downregulation in fibroblasts leads to cytosolic sensing of DNA and interferon signaling induction. (A) ISG mRNA expression, represented as an ISG score (i.e. the median fold change of mRNA levels of seven ISGs (*RSAD2, OAS1, Mx1, IFI27, ISG15, IRF7, IFI44L*), and interferon beta (*IFNb*) mRNA expression in control and ATAD3A patient-derived primary fibroblasts, vehicle-treated (DMSO) or treated with 300 μM 4,4′-diisothiocyanostilbene-2,2′-disulfonic acid (DIDS) or 100 nM rapamycin for 72 hours. Mean values and data points of five to seven independent experiments are shown. * indicates statistical significance in 2-way ANOVA with Dunnett’s multiple comparison tests between treatment conditions. # indicates statistical significance in 2-way ANOVA with Dunnett’s multiple comparison test between control and patient-derived fibroblasts for each treatment. (B) Representative confocal images of control (HDF 1, 2, 3) and *ATAD3A* patient-derived primary fibroblasts (AGS1530, AGS2216, AGS2792) stained for the nucleus (NucBlue in blue) and mitochondria (Mitotracker in yellow). Scale bar = 50 μm. (C) Mitochondrial morphology quantification of control and ATAD3A fibroblasts was performed via supervised machine learning (ML) training using HDF 1 cells with fragmented (CCCP-treated), normal (DMSO-treated), and hypertubular (CHX-treated) mitochondria. Automatic single-cell trinary classification of control (HDF 1, 2, 3) and *ATAD3A* patient fibroblasts by supervised ML. Data represent mean ± SD of 3 independent experiments, (200-442 cells per cell line), non significant in one-way ANOVA. (D) Representative images of immunofluorescence staining of mitochondria (Mitotracker, magenta), double stranded DNA (DNA, green) and nucleus (DAPI, blue) in BJ-5ta fibroblasts knocked-down for ATAD3A by 2 siRNA (siATAD3A 1 and 2), observed by confocal microscopy. Mitochondria/DNA overlap is represented in white. Bottom panel shows a magnification of indicated frames in top panel. Scale bar represents 10 μm in top panel and 5 μm in bottom panel. Graph on the right shows a quantification of DNA spots not colocalized with mitochondrial staining per cell. Means are symbolized by red bars. Results averaged from at least 25 cells counted from one experiment representative of three independent experiments. **** indicates statistical significance in one-way ANOVA with Tukey’s multiple comparison test. ns: non significant.

We report mutations in *ATAD3A* to be associated with an up-regulation of type I interferon signaling equivalent to that seen in established type I interferonopathies. The identification of asymptomatic individuals heterozygous for *ATAD3A* null alleles (Harel et al., 2016), and the results of in vitro functional testing relating to pathogenic missense mutations, including the substitutions recorded in our patients (Cooper et al., 2017; Harel et al., 2016, this work), indicate that such missense mutations act in a dominant-negative fashion, likely through interfering with normal oligomer function (Gilquin et al., 2010). Although we cannot rule out the possibility of some degree of interferon signaling mediated by RNA, knockdown of *ATAD3A* in THP-1 cells induced type I interferon signaling which was dependent on the cGAS-STING dsDNA sensing pathway. Concordantly, removal of mtDNA abrogated this effect. Both findings were also observed in patient fibroblasts. Mitochondrial dysfunction and altered mitochondrial ultrastructure have been reported to cause mtDNA leakage into the cytosol (Kim et al., 2019). Thus, one mechanism to explain the observed cGAS-STING-mediated induction of interferon would be through cytoplasm sensing of mtDNA. In mammalian cells, ATAD3A is a mitochondrial membrane protein enriched at contact sites between mitochondria and the endoplasmic reticulum (Baudier, 2018). We did not observe significant changes in mitochondrial protein (VDAC1, TOM20) levels in THP-1 cells or patient-derived fibroblasts (Fig. 2 B, Fig. 3 C, Fig. 4 E, Fig. S2 D, Fig. S3 B), suggesting that mitochondrial mass per se is unaffected by ATAD3A dysfunction. Cooper et al. recorded fragmented mitochondria on overexpression of the p.(Gly355Asp) mutation in fibroblasts, and in neurons derived from patient fibroblasts with this substitution (Cooper et al., 2017). However, in keeping with a previous study (Dorison et al., 2020), we did not observe any abnormalities in fibroblasts from Patients 2, 3 and 4 on quantification of mitochondrial network shape using a supervised machine learning approach (Fig. 5, B and C), or in *ATAD3A*-silenced fibroblasts (Fig. 5 D). Aggregated/larger mtDNA foci have been described in cells of patients with *ATAD3A* deletions (Desai et al., 2017). Downregulation of TFAM, the mtDNA nucleoid organizer which ATAD3A binds (He et al., 2012), also causes enlarged mtDNA foci and leads to cytosolic leakage of mtDNA (West et al., 2015). We found total TFAM protein levels to be similar in patient and control cells (Fig. 4 E), ruling out TFAM loss as a potential mechanism of increased mtDNA sensing. However, mutated ATAD3A could alter the retention of mtDNA by TFAM, as reported when ATAD3A expression and oligomerization are altered (Zhao et al., 2019). Consistent with this, we observed increased numbers of cytosolic DNA foci upon ATAD3A downregulation and a contribution of VDAC1 channels to enhanced interferon signaling which is worthy of more detailed study. Complementarily to its action as an anti-inflammatory drug (Wang et al., 2020), the effect of rapamycin in patient fibroblasts suggests that it might act on a cell intrinsic basis, possibly inducing the autophagy of cytosolic DNA or of damaged mitochondria leaking mtDNA, to curb mtDNA detection and interferon signaling. Of note, the mitophagy-inducing potential of rapamycin has been implicated in therapeutic strategies relating to mitochondrial disease (Li et al., 2014; Villanueva Paz et al., 2016).

Two of the patients that we ascertained demonstrate signs consistent with standardized criteria for a diagnosis of the autoimmune disorder systemic sclerosis (van den Hoogen et al., 2013), with classical histological features of systemic sclerosis (i.e. fibrosis, loss of skin adnexae and deep lymphocytic peri-vascular infiltrates) on skin biopsy in both cases. The positivity for multiple autoantibodies, and elevated levels of gamma-globulin observed in Patient 2 are suggestive of an autoimmune phenomenon, although these features were not present in Patient 3. The pathology of systemic sclerosis remains unclear, with the phenotype likely representing the end point of a heterogeneous set of disease processes (Bhattacharyya et al., 2011). Genetic association (Lopez-Isac et al., 2019), gene expression (Brkic et al., 2016) and experimental studies (Ah Kioon et al., 2018; Wu et al., 2019) indicate a link between systemic sclerosis and up-regulated type I interferon signaling, presumed to be induced by self-derived DNA or RNA (Barrat et al., 2016). The source of such nucleic acids remains unclear. While the potential of mtDNA to induce cytoplasmic cGAS-STING-mediated interferon signaling (West et al., 2015; West and Shadel, 2017), and thereby promote autoimmunity (Kim et al., 2019), has been recently highlighted (Crow, 2019), it is possible that other inflammatory signaling pathways might also be relevant (Zhou et al., 2011; Yu et al., 2020).

Although mutations in ATAD3A are associated with a disturbance of mitochondrial structure, oxidative phosphorylation enzyme deficiency has been only variably recorded in affected individuals (Desai et al., 2017; Gunning et al., 2020; Harel et al., 2016; Peralta et al., 2019; Frazier et al., 2020), indicating the possibility of disease-related pathology due to other than a failure of energy production. Whilst the recurrent observation of absence of the eyelashes and eyebrows, seen in four of seven patients in our series, suggests a developmental aspect to the function of ATAD3A, our findings indicate that type I interferon induction might also represent a pathogenic factor. In this regard, we note that intracranial calcification, a sign observed in a number of type I interferonopathies, was present in Patient 6; and that dystonia, peripheral neuropathy and hypertrophic cardiomyopathy, as recorded in some patients with mutations in *ATAD3A*, are (also) features of interferon-related disease (Crow et al., 2015). Additionally, isolated spastic paraparesis (as seen in Patient 7) is a well-recognized manifestation of mutations in several type I interferonopathy related genes (Crow et al., 2014).

Our clinical data are important in extending the phenotype associated with mutations in *ATAD3A* to include an association with systemic sclerosis. Further, we note the marked difference in phenotype between the mother (Patient 7 here) (lower limb spastic paraparesis with normal intellect) and her son (four-limb spastic dyskinesia and severe intellectual delay) as reported by Cooper et al. (Cooper et al., 2017). Variability in clinical expression, including purely neurological disease (Livingston et al., 2014), frank autoimmunity (Van Eyck et al., 2015) and complete clinical non-penetrance (Lepelley et al., 2020; Rice et al., 2020), is highly characteristic of the type I interferonopathies.

To our knowledge, mutations in *ATAD3A* represent the first example of a Mendelian mitochondrial disease directly implicating mtDNA in type I interferon induction. The relative contribution of enhanced type I interferon signaling to the ATAD3A-associated phenotype is yet to be determined; however, the advent of strategies to block interferon signaling may allow this question to be tested in an experimental medicine setting in the near future.

## Materials and methods

### Samples obtained from patients

Samples were obtained from the probands and parents with written informed consent. The study was approved by the Comité de Protection des Personnes (ID-RCB / EUDRACT: 2014-A01017-40) and the Leeds (East) Research Ethics Committee (10/H1307/132).

### Histological analyses

Formalin-fixed paraffin-embedded skin biopsy slides were analyzed after hematoxylin and eosin (H&E) staining.

### Genetic studies

DNA was extracted from whole blood using standard methods. Exome sequencing was performed on genomic DNA from Patients 1 and 2 and their parents using a SureSelect Human All Exon kit (Agilent Technologies) for targeted enrichment, and Illumina HiSeq2000 for sequencing. Variants were assessed using the in silico programs SIFT (http://sift.jcvi.org) and Polyphen2 (http://genetics.bwh.harvard.edu/pph2/). Population allele frequencies were obtained from the gnomAD database (http://gnomad.broadinstitute.org). Sanger sequencing was performed to confirm the identified *ATAD3A* variants. The reference sequence used for primer design and nucleotide numbering was NM_001170535.

### Interferon status

Whole blood was collected into PAXgene tubes (Qiagen) and total RNA extracted using a PreAnalytix RNA isolation kit. Interferon scores were generated in one of two ways; as previously described using TaqMan probes to measure the mRNA expression of six interferon-stimulated genes (*IFI27, IFI44L, IFIT1, ISG15, RSAD2, SIGLEC1*) normalized to the expression level of *HPRT1* and *18S rRNA* (Rice et al., 2013). The median fold change of the interferon-stimulated genes is compared to the median of 29 healthy controls to create an interferon score for each individual, with an abnormal score being defined as greater than 2.46. For NanoString ISG analysis, total RNA was similarly extracted from whole blood with a PAXgene (PreAnalytix) RNA isolation kit. Analysis of 24 genes and 3 housekeeping genes (table immediately below) was conducted using the NanoString customer designed CodeSets according to the manufacturer’s recommendations (NanoString Technologies, Seattle, WA). Agilent Tapestation was used to assess the quality of the RNA. 100ng of total RNA was loaded for each sample. Data were processed with nSolver software (NanoString Technologies Seattle, WA). The data was normalized relative to the internal positive and negative calibrators, the 3 reference probes and the control samples. The median of the 24 probes for each of 27 healthy control samples was calculated. The mean NanoString score of the 27 healthy controls +2SD of the mean was calculated. Scores above this value (2.75) were designated as positive.

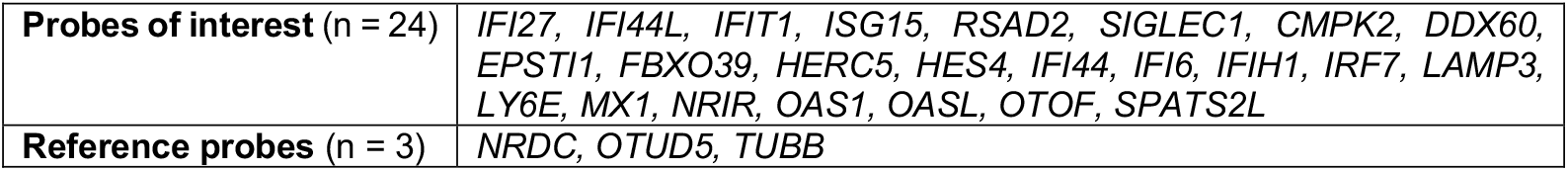

Plasma and cerebrospinal fluid interferon alpha (IFN-α) concentrations (fg/ml) were determined by single molecule array (Simoa) using a commercial kit for IFN-α2 quantification (Quanterix™, Lexington, MA, USA). The assay was based on a three-step protocol on an SR-X analyzer (Quanterix).

### Cell culture

Human embryonic kidney (HEK) 293FT cells (catalog no. R70007; Invitrogen) were grown in DMEM (GIBCO) supplemented with 10% (v/v) fetal bovine serum (GIBCO). Primary fibroblasts (3 from controls and from patients 2 (AGS1530), 3 (AGS2216) and 4 (AGS2792)) and BJ-5ta (catalog no. CRL-4001; ATCC) fibroblasts were grown in DMEM supplemented with 10% (v/v) fetal bovine serum, with 1% penicillin-streptomycin and 0.25 μg/mL amphotericin B (GIBCO) added for primary fibroblasts. THP-1 cells were from ATCC (TIB-202) and wild-type (WT), STING knockout (KO) and cGAS KO THP-1 cells were from InvivoGen (THP-1 Dual, ref. thpd-kostg and thpd-kocgas). MAVS KO THP-1 cell lines were recently generated in the Rehwinkel laboratory (Hertzog et al., 2020 *Preprint*). All THP-1 cell lines were maintained in RPMI (Invitrogen) supplemented with 10% (v/v) fetal bovine serum, 10 mM HEPES and 0.05 mM 2-mercaptoethanol (GIBCO). U2OS cells (catalog no. HTB-96; ATCC) were maintained in McCoy medium (Invitrogen) supplemented with 10% (v/v) fetal bovine serum. All cell lines were cultured at 37°C in 5% CO_2_. Primary fibroblasts were treated for 72 hours with either vehicle (DMSO, Sigma-Aldrich), 300 μM 4,4′-diisothiocyanostilbene-2,2′-disulfonic acid (DIDS, Sigma-Aldrich 309795), 100 nM rapamycin (Selleckchem S1039) or 50-200 μM of BAX inhibitor peptide V5 (MedChemExpress HY-P0081). Peripheral blood mononuclear cells (PBMCs) were isolated by Ficoll-Paque density gradient (Lymphoprep, Proteogenix) from the blood of patients and healthy donors. Control PBMCs from healthy donors were obtained from the Etablissement Français du Sang blood bank. Cells were pelleted and kept at -80C° until further analysis.

### Knockdown of gene expression by short hairpin RNA (shRNA)

Lentiviral constructs containing short hairpin RNA (shRNA) targeting *ATAD3A* (TRCN0000136391: shATAD3A 91, TRCN0000135329: shATAD3A 29, TRCN0000136793: shATAD3A 93), *TFAM* (TRCN0000329820: shTFAM) and control Luciferase (shLuc, SHC007) were made by ligating annealed oligonucleotides into pLKO.1 (TRC cloning vector, a gift from David Root (Addgene plasmid #10878)), according to the RNAi consortium protocol (https://portals.broadinstitute.org/gpp/public/). pLKO.1 empty vector was from Open Biosystems, and pLKO.1 control shRNA (scramble shSCR, SHC002 and non target shNTgt, SHC016) from Sigma-Aldrich. Lentiviral vectors carrying these constructs were produced by calcium phosphate transfection of 293FT cells with shRNA constructs in combination with packaging vectors psPAX2, a gift from Didier Trono (Addgene plasmid #12260), and envelope pCMV-VSV-G (Addgene plasmid #8454). Medium of 70% confluent 293FT in 75 cm^2^ flasks was changed 2 hours before transfection. Calcium phosphate precipitates were prepared by mixing 12.5 μg shRNA vectors with 12.5 μg psPAX2 and 5 μg pCMV-VSV-G in water for a final volume of 875 μL. 125 μL 2 M CaCl_2_ and 1 mL HBS 2X (50 mM HEPES, 10 mM KCl, 280 mM NaCl, 1.5 mM Na_2_HPO_4_, pH 7.05) were sequentially added dropwise in slowly vortexed solution. Solutions were incubated at RT for 20 minutes and mixed gently with 293FT supernatant. Medium was replaced by 7 mL of culture medium 24 hours later. After 24 more hours, supernatants were collected, centrifuged at 1,700 rpm for 5 minutes and 0.45 µm-filtered. 500,000 THP-1 cells were transduced with 0.5 mL lentiviral vectors, 8 μg/mL polybrene (Millipore) and 10 mM HEPES (Invitrogen) in 12-well plates and medium replaced 24 hours later. Two days after transduction, transduced cells were selected with 0.5-8 μg/mL puromycin (Sigma-Aldrich). Cells were collected for analysis four days after transduction.

### Knockdown of gene expression by CRISPR

Single guide RNAs (sgRNA) targeting *ATAD3A* (oligonucleotides used given in table below) were designed using the CRISPOR tool (crispor.tefor.net) and cloned into lentiviral construct lentiCRISPRv2-hygro, a gift from Brett Stringer (Addgene plasmid #98291) (Stringer et al., 2019), following the protocol provided on the plasmid website. Empty vectors #3 and #12 are two clones obtained after BSMBI digestion, to remove buffer sequence, blunting and ligation using the Quick blunting kit (NEB), following the manufacturer’s instructions. Lentiviral particles were produced, and THP-1 cells transduced as described for shRNA. Two days after transduction transduced cells were selected with 500 μg/mL hygromycin B (Invivogen), and selected cell pools analyzed seven days after transduction.

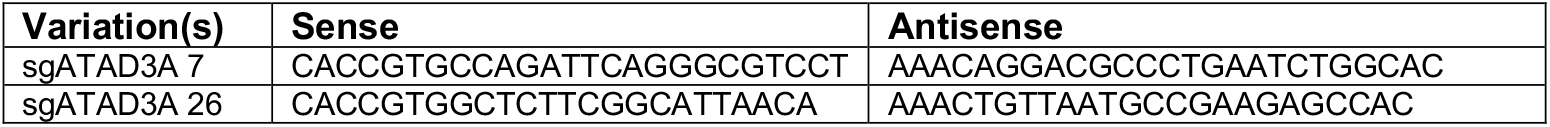

### Knockdown of gene expression by small interfering RNA (siRNA)

Expression of *cGAS, MAVS, BAX, BAK1, ATAD3A* and *TFAM* in control or patient-derived primary fibroblasts was downregulated by using 20 nM of siRNA (references in the table below). Negative control or gene-targeting siRNAs were delivered to cells using the Lipofectamine RNAiMAX Transfection Reagent (Invitrogen) according to the manufacturer’s instructions. Cells were analyzed 72 hours after transfection.

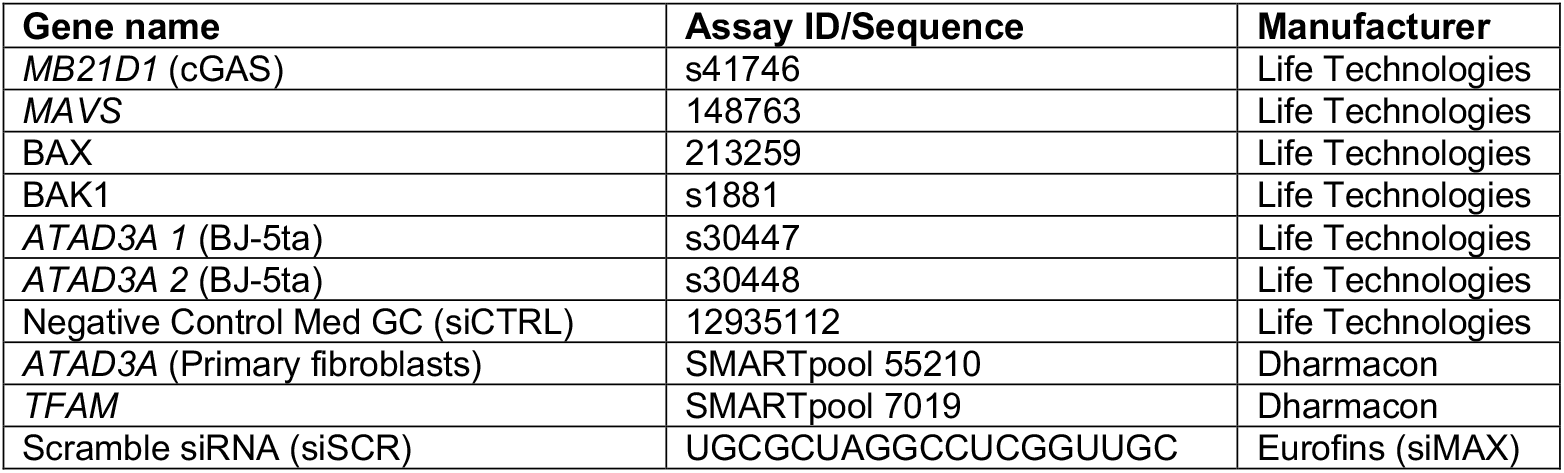

### Mitochondrial DNA depletion

mtDNA depletion in THP-1 cells was induced by low dose ethidium bromide (100 ng/mL) treatment in medium supplemented with 50 μg/mL uridine (Sigma-Aldrich) and 1 mM sodium pyruvate (GIBCO) for seven to fourteen days before shRNA knockdown. In primary fibroblasts, mtDNA depletion was induced by 100 μM 2′,3′ dideoxycytidine (ddC, Sigma-Aldrich) in medium supplemented with uridine and sodium pyruvate, as for THP-1 cells, for seven to fourteen days before analysis. To control for mtDNA depletion, total DNA from 0.5.10^6^ cells was extracted using the DNeasy Blood and Tissue Kit (Qiagen), following the manufacturer’s instructions. DNA concentrations were determined by photometry (Nanodrop) and 15 ng, 7.5 ng and 3.75 ng DNA were used to perform qPCR for the mitochondrial gene *MT-COXII* and the nuclear gene *GAPDH* (primers given in table below). Quantitative PCR (qPCR) was performed using Power SYBR Green (Invitrogen). Ratios of 2^-CT^ for *MT-COXII* over *GAPDH* for the different DNA concentrations were averaged and fold of untreated (UT) is shown in the figures.

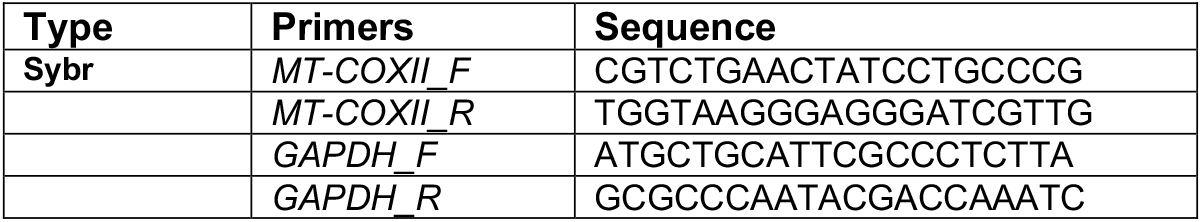

### Generation of U2OS cells stably expressing STING

The pMSCV-hygro plasmid carrying WT *STING1* cDNA (Addgene #102598) was used in combination with packaging vector pCL-Ampho (Novus) and envelope vector pCMV-VSV-G (Addgene plasmid #8454) to produce retroviral vectors as described for shRNA lentiviral vectors. 100,000 U2OS cells were transduced with 0.5 mL retroviral vectors, 8 μg/mL polybrene (Millipore) and 10 mM HEPES (Invitrogen) in 12-well plates and medium replaced 24 hours later. Two days after transduction, transduced cells were selected and maintained in culture with 200 μg/mL hygromycin (Invivogen). STING expression was verified by western blotting.

### Overexpression of WT and mutant ATAD3A

pCMV6-Entry vector encoding Myc-DKK-tagged human wild type (WT) ATAD3A cDNA (NM_018188.4, Q9NV17-1) was obtained from Origene and mutations were inserted by site-directed mutagenesis using the Q5 kit (E0554S; New England Biolabs NEB) according to the manufacturer’s instructions and using primers in the table below. To obtain plasmids encoding the most common ATAD3A isoform (NM_001170535.2, Q9NV17-2), the sequence comprising the beginning of exon 3 was removed using the Q5 kit and primers below. pCMV6-Myc-DKK empty vector (EV) was generated by digestion with EcoRI and MluI restriction enzymes (ThermoFischer Scientific), end blunting (Quick blunting kit; NEB), and ligation with (T4 ligase, NEB), following the manufacturer’s instructions. Constructs were verified by Sanger sequencing.

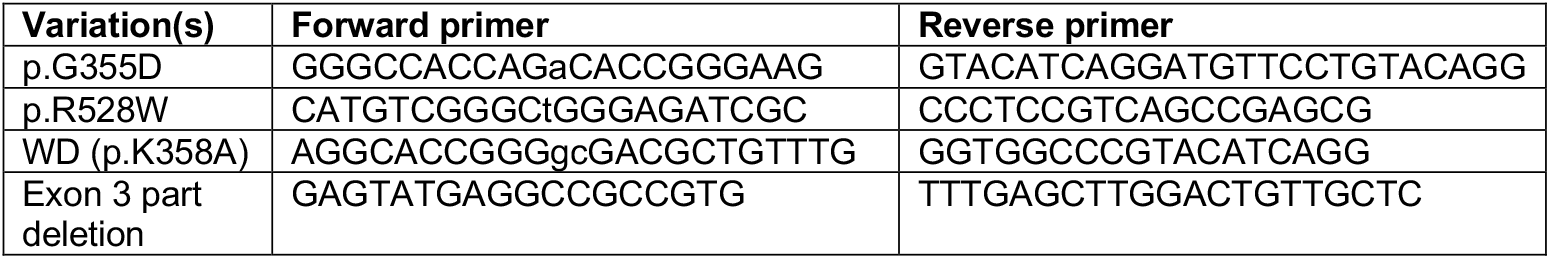

At 70% confluency, U2OS-STING cells plated in 12-well plates were transiently transfected using TransIT®-2020 (Mirus) with 500 ng of ATAD3A WT, G355D, R528W or WD cDNA in pCMV6-Myc-DDK plasmid. 5 hours after transfection, medium was changed and 24 hours after transfection, cells were treated with 10 μM Q-VD-OPh (Sigma-Aldrich SML0063). Cells were collected 24 hours later for RNA and protein analysis.

### SDS-PAGE and western blot analysis

For whole cell lysate analysis, proteins were extracted from THP-1 cells, PBMCs and primary fibroblasts using lysis buffer (RIPA, 1% protease inhibitor, 1% phosphatase inhibitor). Bolt LDS Sample Buffer (4X) (Novex Life Technologies) and Bolt Sample Reducing agent (10X) (Novex Life Technologies) were added to protein lysates, samples resolved on 8% or 4-12 % Bis-Tris Plus gels (Invitrogen) and then transferred to nitrocellulose membrane for 7 min at 20 V using the iBlot 2 Dry Blotting System (Invitrogen). Where protein phosphorylation status was investigated, membranes were blocked in LI-COR® buffer, and primary phospho-antibodies incubated for 48 hours in the blocking solution. Otherwise, membranes were blocked with 5% non-fat milk in TBS, and primary antibodies incubated overnight at 4°C in 1.5% Bovine Serum Albumin in TBS buffer supplemented with 0.1% Tween for most antibodies; in 2.5% non-fat milk in TBS buffer supplemented with 0.1% Tween for anti-STING antibody; and in 1.5% Bovine Serum Albumin in TBS buffer supplemented with 0.4% Tween for rabbit anti-ATAD3 antibodies. A list of antibodies used in this study is supplied in table below. Membranes were washed and incubated with appropriate anti-mouse or anti-rabbit secondary antibodies for 45 minutes at room temperature (LI-COR® System). Signal was detected using the Odyssey®CLx System (LI-COR®). Comparative signal analyses were performed using Fiji (ImageJ).

| Antibodies | Host species | Source | Catalog reference |
| --- | --- | --- | --- |
| Cofilin (loading control) | Rabbit monoclonal | Cell Signaling | #5175 |
| β-Actin (loading control) | Rabbit monoclonal | Cell Signaling | #3700 |
| Vinculin (loading control) | Mouse monoclonal | Santa Cruz | sc-73614 |
| ATAD3 | Mouse monoclonal | Santa Cruz | sc-376185 |
| ATAD3 | Rabbit polyclonal | Gift of J. Baudier | N-Ter (Gilquin et al., 2010) |
| IRF3 | Rabbit monoclonal | Cell Signaling | #11904 |
| Phospho-(Ser396)IRF3 | Rabbit monoclonal | Cell Signaling | #4947 |
| STING | Mouse monoclonal | R&D Systems | MAB7169 |
| Phospho-(Ser366)STING | Rabbit monoclonal | Cell Signaling | #19781S |
| TFAM | Mouse monoclonal | Santa Cruz | sc-376672 |
| TOM20 | Mouse monoclonal | Santa Cruz | sc-17764 |
| VDAC1 | Mouse monoclonal | Santa Cruz | sc-390996 |
| ISG15 | Rabbit polyclonal | Abcam | ab14374 |
| MX1 | Rabbit polyclonal | Abcam | ab95926 |
| BAX | Rabbit polyclonal | GeneTex | GTX109683 |
| BAK1 | Rabbit polyclonal | GeneTex | GTX100063 |
| FLAG (M2) | Mouse monoclonal | Signam-Aldrich | F3165 |
| IRDye 800CW<br>(secondary antibody) | Goat anti-mouse | Li-COR | 926-32210 |
| IRDye 800CW<br>(secondary antibody) | Goat anti-rabbit | Li-COR | 926-32211 |
| IRDye 640RD<br>(secondary antibody) | Goat anti-mouse | Li-COR | 926-68070 |
| IRDye 640RD<br>(secondary antibody) | Goat anti-rabbit | Li-COR | 926-68071 |

### RT-qPCR quantification of gene expression

Total RNA was extracted using the RNAqueous-Micro Kit (Ambio), and reverse transcription performed with the High-Capacity cDNA Reverse Transcription Kit (Applied Biosystems). Levels of cDNA were quantified by RT-qPCR using Taqman Gene Expression Assay (Applied Biosystems). Differences in cDNA inputs were corrected by normalization to *HPRT1* cDNA levels. Relative quantitation of target cDNA was determined by the formula 2^-ΔCT^, with ΔCT denoting fold increases above control. When indicated, an ISG score was calculated as the median of the fold change expression of *RSAD2, OAS1, Mx1, IFI27, ISG15, IRF7, IFI44L*, unless otherwise stated, and used as a readout. A list of the probes used in this study is supplied in the table below.

| Type | Primers | Assay ID |
| --- | --- | --- |
| Taqman | *HPRT1* | Hs03929096_g1 |
|  | *IFI27* | Hs01086370_m1 |
|  | *IFI44L* | Hs00199115_m1 |
|  | *IFIT1* | Hs00356631_g1 |
|  | *ISG15* | Hs00192713_m1 |
|  | *RSAD2* | Hs01057264_m1 |
|  | *SIGLEC1* | Hs00988063_m1 |
|  | *RSAD2* | Hs01057264_m1 |
|  | *OAS1* | Hs00973637_m1 |
|  | *Mx1* | Hs00895608_m1 |
|  | *IRF7* | Hs01014809_g1 |
|  | *IFNB1* | Hs01077958_s1 |
|  | *IFNA2* | Hs00265051-s1 |
|  | *IFNG* | Hs00989291-m1 |
|  | *TFAM* | Hs00273372_s1 |
|  | *ATAD3A* | Hs01587333_m1 |
|  | *MB21D1 (cGAS)* | Hs00403553_m1 |
|  | *MAVS* | Hs00920075_m1 |
|  | *BAK1* | Hs00832876_g1 |

### DNA immunofluorescence staining and confocal microscopy

BJ-5ta cells were grown on a glass coverslip in a 6-well plate 24 hours prior to treatment with indicated siRNAs. Three days post siRNA treatment, cells were labelled with Mitotracker Red CMXRos (100 nM) for 30 minutes. Cells were washed and fixed with 4% paraformaldehyde in phosphate buffered saline (PBS) and permeabilized with 0.25% Triton X-100, followed by blocking in 3% bovine serum albumin/PBS for 1 hour. Cells were incubated with primary antibody (mouse monoclonal anti-DNA (AC-30-10, Progene) used at 1:200 concentration) overnight in a humidified chamber at 4°C followed by extensive washing before addition of secondary antibody (goat anti-mouse IgG, IgM (H+L), Alexa 488 at 1:300 concentration) for 1 hour at room temperature. Slides were mounted in Vectashield antifade aqueous mounting medium (Vector Laboratories), with 4′,6-diamidino-2-phenylindole (DAPI). Single plane images were unbiasedly acquired with Nikon A1R+ confocal microscope using a 60.0X / 1.4 oil objective. Analysis was performed on single focal planes acquired. Images were processed and analyzed using ImageJ Fiji. Since the mitochondrial networks were similar between conditions, the number per cell of DNA-stained foci not overlayed with the mitochondrial stain, and thus cytosolic, was counted. 3 to 8 fields of view were analyzed per condition per experiment in a blinded fashion.

### Automated imaging of mitochondrial morphology

Human fibroblasts cells were seeded on CellCarrier-96well Ultra microplate (Perkin Elmer) and incubated for at least 24 hours in growth media. Nuclei were labeled with NucBlue™ Live ReadyProbes™ Reagent (ThermoFisher Scientific). Fluorescent labeling of mitochondria was achieved using Tetramethylrhodamine Ethyl Ester Perchlorate (TMRE) and MitoTracker DeepRed at 100 nM for 30 minutes at 37°C, 5% CO_2_. Cells were treated with 5 μM protonophore carbonyl cyanide m-chlorophenyl hydrazone (CCCP, Sigma-Aldrich) or 10 μM cytosolic protein synthesis inhibitor cycloheximide (CHX, Sigma-Aldrich) to induce mitochondrial fragmentation and hypertubulation, respectively. Spinning disc confocal images were acquired using the Operetta CLS High-Content Analysis systems (Perkin Elmer), with 63x Water/1.15 NA. TMRE (530-560 nm), MitoTracker DeepRed (615-645nm), and DAPI (355-385 nm) were excited the appropriate LEDs (Operetta CLS). Harmony Analysis Software (PerkinElmer) was used for automated confocal image analysis as described in detail in Cretin et al. (2021). Z-projected images first undergo brightfield correction. Nuclei and cellular segmentation were defined using the “Find Nuclei” building block with the HOECHST 33342 channel and the “Find Cytoplasm” building block with the Alexa 633 Mitotracker DeepRed (MTDR, mitochondria) channel. Mitochondrial network was analyzed using SER Texture properties (Ridge, Bright, Edge, Dark, Valley, Spot) and the PhenoLOGIC supervised machine learning algorithm was used to identify the best properties able to segregate the three populations: “Normal”, “Fragmented” and “Hypertubulated” network. 81-104 cells of each control (Normal: WT + DMSO at t=0, Fragmented: WT + CCCP at t=3h, Hypertubulated: WT + CHX at t=6h) were selected to feed the algorithm for training. Automatic single-cell classification of non-training samples (i.e. unknowns) was carried out by the supervised machine-learning module.

### Quantification and statistical analysis

Statistical analysis was conducted using GraphPad Prism 8 software. Data were tested for normal distribution of variables and displayed as means ± standard error of the mean (sem) unless otherwise noted. Measurements between two groups were performed with a Student’s t test if normally distributed, or Mann-Whitney test otherwise. Groups of three or more were analyzed by one-way analysis of variance (ANOVA) or the Kruskal-Wallis test, or paired ANOVA or Friedman test. Grouped analyses were interrogated by two-way ANOVA with post-hoc multiple comparison tests. Values of n repeats and statistical parameters for each experiment are reported in the figures and figure legends. p < 0.05 was considered significant.

### Clinical reports

#### Patient 1

This female was born at term following a normal pregnancy. She demonstrated excessive sleepiness in the first two weeks of life with feeding difficulties. Developmental delay was obvious by age five months. Neopterin was raised (256 nmol/l – normal, 7 -65 nmol/l) in CSF at age two years. She had extremely blond hair (unlike her parents or half-siblings), absent eyelashes and eyebrows, and deep-set eyes. She was hypotonic centrally and subsequently exhibited profound global developmental delay, visual impairment and variable dystonia. She developed seizures at five years of age which have been difficult to manage, requiring ventilation on two occasions for status epilepticus. At the age of nine years she demonstrates profound delay, rigid dystonia and no useful motor function. Communication is limited to eye movements and facial expression. Her head circumference is on the 50^th^ centile, and cranial MRI showed some high signal in the anterior temporal lobes and at the parietal ventricular poles, with no obvious calcification.

#### Patient 2

This female was noted to have an absence of the eyelashes and eyebrows, myopia, and early-onset alacrima in the first year of life. She required corneal grafts at age five years because of bilateral keratitis secondary to her alacrima. She subsequently developed features of autoimmunity including antibody-positive Hasimoto’s thyroiditis and vitiligo, also demonstrating growth hormone deficiency. Mild lower limb spasticity was noted in infancy, prompting cerebral MRI at age 6 years and a CT one year later, which were both normal. She attended university, with no intellectual into adulthood, but demonstrates a spastic diplegic gait which appears stable. Beginning at 21 years of age she developed sclerodermatous involvement of the hands, face and ventral surface of the forearms, which progressed rapidly over a few months. When reviewed aged 22 years she exhibited small joint swelling and diffuse skin thickening, particularly involving the fingers of both hands and extending proximal to the metacarpophalangeal joints (Rodnan skin score: 33/51), fulfilling ACR/EULAR) criteria for a diagnosis of systemic sclerosis. There was an absence of digital ulcerations. Immunoglobulin levels, antinuclear (1/5220) and anti-dsDNA antibody titers were persistently elevated. Systemic sclerosis-related antibodies were not detected. Skin biopsy of affected tissue showed classical features of systemic sclerosis, with massive collagen fibrosis, loss of elastic fibers and adnexal epithelia, and deep peri-vascular lymphocytic infiltrates in the absence of immunoglobulin or complement deposition. In view of the rapid clinical progression, oral corticosteroids were initiated, resulting in a prompt remission of arthralgia and joint swelling. However, given no apparent effect on skin thickening and hardening after 11 months of treatment, rapamycin was introduced, and was associated with a partial regression of the cutaneous fibrosis and a decrease of the Rodnan skin score to 10/51 over the following two years.

#### Patient 3

This female demonstrated early onset severe motor delay with impaired cognition but good social contact. Extensive genetic investigations were negative, including array comparative genomic hybridization. Within the first year of life she developed feeding difficulties, with relatively rapid feeding noted to easily promote vomiting, eventually necessitating the placement of a gastrostomy at three years of age. Respiratory chain enzyme analysis in fibroblasts was normal (Bird et al., 2019). From age two years she developed a progressive hardening of the skin, and a change in facial appearance due to loss of fat tissue, and areas of panniculitis involving the lower limbs. At four years of age she exhibited peri-oral sclerosis, hard and tight sclerodermatous skin of the upper and lower limbs, and severe flexion contractures at the knees, ankles, elbows, wrists and fingers (Rodnan skin score: 32/51). Skin biopsy revealed deep dermal sclerosis and loss of skin appendages, thick collagen bundles and mild lymphocytic infiltration. MRI imaging of the upper limbs confirmed cutaneous thickening. Treatment with rapamycin and low dose corticosteroids was initiated at age four years, leading to resolution of nightly unrest and cessation of pain medication, reduced induration of sclerotic limb lesions, improved mouth closure and the reintroduction of supplementary oral feeding.

#### Patient 4

This male, now aged two years, demonstrates severe developmental delay with central hypotonia, peripheral dystonia, a peripheral neuropathy and hypertrophic cardiomyopathy. There was a deficit of respiratory chain complex I and V on muscle biopsy, and a lactate peak seen on cerebral MRI, both before the age of 1 year. A cranial CT at age two years did not demonstrate any evidence of intracranial calcification.

#### Patient 5

This seven year old male (patient II-I, Family 3 in (Harel et al., 2016)) exhibits global developmental delay, with truncal hypotonia and peripheral spasticity. There is an absence of the eyebrows, optic nerve pallor, and macular hypoplasia. An echocardiogram revealed hypertrophic cardiomyopathy, initially detected at age three months. Electrophysiology identified a peripheral neuropathy. Additionally, he has growth hormone deficiency and demonstrates extensive vitiligo. Plasma lactate was intermittently elevated, and a lactate peak was seen on MR spectroscopy.

#### Patient 6

This two year old female of Iranian ancestry (described in (Hanes et al., 2020)) required the removal of bilateral cataracts at age three months. At five months old she demonstrated axial hypotonia and hyporeflexia. Neurophysiology defined a sensorimotor polyneuropathy with axonal features which was confirmed on nerve and muscle biopsy. By eight months of age developmental regression became apparent, so that she was no longer able to sit unsupported and she had stopped rolling. Cranial MRI was unremarkable at this time. She was diagnosed with profound bilateral sensorineural hearing loss. At age 18 months she presented with bilateral ophthalmoplegia, partial left-sided ptosis and visual impairment, as well as upper and lower limb dystonic posturing that did not respond to levodopa-carbodopa. At 21 months of age she developed refractory epilepsy, and at age 24 months a tracheostomy was placed due to intermittent central and obstructive apneas and an inability to wean off mechanical ventilation. Repeat MRI showed progressive cerebral and cerebellar atrophy between studies performed at 20 and 24 months of age. MR spectroscopy and repeated serum lactate measurements were normal. At age 32 months she remains ventilated via tracheostomy with little interaction with her surroundings and ongoing seizures and dystonia.

#### Patient 7

This patient, also reported previously (Cooper et al., 2017), was noted to demonstrate bilateral lower limb spasticity resulting in Achilles tendon lengthening at an early age. Now, aged 40 years, she exhibits severe spastic paraparesis. She is of normal intellect and performed well academically, her only other problem being myopia (minus 10 diopters) with significant photophobia resulting in restricted eye opening. Her son, also reported by Cooper et al. (Cooper et al., 2017), but not tested here, was noted at age four months to exhibit lower limb spasticity. Spinal and cranial MRI at 12 months of age were normal. At the age of three years he demonstrated dyskinetic cerebral palsy with severe upper and lower limb spasticity, was unable to speak, sit or stand, but could crawl by pulling himself forwards with his arms and roll on the floor. Similar to his mother, there is an absence of the lateral eyelashes and photophobia.

## Abbreviations

ATAD3A: ATPase family AAA domain-containing protein 3A
CSF: cerebrospinal fluid
CT: computed tomography
cGAS: cyclic GMP-AMP synthase
ds: double stranded
HDF: human dermal fibroblasts
IFNb: interferon beta
IRF3: Interferon regulatory factor 3
ISG: interferon-stimulated gene
MAVS: mitochondrial-antiviral signaling protein
MDA5: melanoma differentiation-associated protein 5
MRI: magnetic resonance imaging
mtDNA: mitochondrial DNA
PBMCs: peripheral blood mononuclear cells
RIG-I: retinoic acid-inducible gene I
shRNA: short hairpin RNA
siRNA: short interfering RNA
STING: stimulator of interferon genes
TFAM: mitochondrial transcription factor A

## Supplemental material summary

Supplementary figure S1 shows clinical, histological and imaging features of systemic sclerosis seen in Patients 2 and 3. Supplementary figure S2 reports interferon induction upon ATAD3A downregulation by CRISPR targeting in THP-1 cells and in patient PBMCs. Supplementary figure S3 reports the study of the signaling pathways involved in interferon induction in patient fibroblasts. Supplementary table 1 summarizes features related to systemic sclerosis in Patients 2 and 3.

## Author contributions

Conceptualization, A.L., B.B.M., C.W., and Y.J.C.; Methodology, A.L., B.B.M., C.W., T.W. and Y.J.C.; Investigation, A.L., E.D.M., E.VN., L.W., S.F., G.I.R., A.D., M.L.F., M.P.R., L.S., E.C., C.B., D.B., B.C. P.dL., L.DS., D.A.D., F.F., L.G., S.H., M.H., H.J.M., M.A.K., K.S., H.T., S.C., T.W., C.W. and B.B.M; Writing – Original draft, Y.J.C.; Writing – Review & editing, A.L., B.B.M., C.W., and Y.J.C.; Funding acquisition, Y.J.C.; Supervision, A.L., and Y.J.C.

## ACKNOWLEDGEMENTS

YJC acknowledges the European Research Council (GA 309449 and 786142 E-T1IFNs), a state subsidy managed by the National Research Agency (France) under the ‘Investments for the Future’ program bearing the reference ANR-10-IAHU-01, and the NIHR UK Rare Genetic Disease Research Consortium. The project was supported by MSDAVENIR (Devo-Decode Project). EVN acknowledges the Research Foundation Flanders (Fonds voor Wetenschappelijk Onderzoek Vlaanderen: grant no. 1S22716N). We sincerely thank Jacques Baudier for the generous use of antibodies. BC is a Senior Clinical Investigator of the Research Foundation-Flanders. Ghent University Hospital, University Hospital Leuven, and Hôpital Universitaire Necker are members of the European Reference Network on Skin disorders (ERN-Skin). This project has received funding from the European Union’s Horizon 2020 research and innovation program under the Marie Skłodowska-Curie grant agreement No.892311 to AL.

