## Supplemental data for "Enhanced cGAS-STING-dependent interferon signaling associated with mutations in ATAD3A"

**Supplementary figure legends**

**Supplementary Figure S1**. **Clinical, histological and imaging features of systemic sclerosis seen in Patients 2 and 3.**

(A) Clinical features of systemic sclerosis seen in Patients 2 and 3. Subpanels a and b highlight sclerodermatous involvement of the hands and legs of Patient 2 at 22 years of age. Similar features were observed in Patient 3 at age 4 years (subpanels e and f). (B) Features of systemic sclerosis seen on skin biopsy. Subpanels c and d, and subpanel g, demonstrate the appearances on skin biopsy observed in Patient 2 and Patient 3 respectively. In subpanel c the overlying epidermis is normal, whilst the dermis is increased in thickness and composed of broad sclerotic collagen bundles extending into the subcutis and replacing fat. The deep dermis shows homogenized collagen and is devoid of elastic tissue. In addition, eccrine glands are very atrophic. The arrectores pilorum of the hair follicles are only recognizable in the upper part of the dermis, because the epithelium is almost completely absent. Subpanel d is from another part of the biopsy, showing the same changes with, in addition, peri-vascular lymphocytic infiltrate in the deep dermis. In subpanel g, similar changes are seen to those observed in Patient 2, but with skin adnexae (hair follicles and sweat glands) still present, consistent with an earlier stage in the same disease spectrum. Scale bars equal 500 µm. (C) MRI of right forearm of Patient 3. Left: TSE T1 sequence before contrast shows diffuse cutaneous thickening in the right forearm, most pronounced on the dorsal-ulnar aspect. Right: TSE T1 sequence after contrast administration (Dotarem 0.5mmol/ml). Arrows indicate signal in the thickened cutis before and after enhancement.

**Supplementary Figure S2**. ***ATAD3A* downregulation by CRISPR editing in THP-1 cells and interferon signalling in ATAD3A patient peripheral blood mononuclear cells (PBMCs).**

(A) qPCR of *ATAD3A* following knockdown of *ATAD3A* by CRISPR editing in THP-1 cell pools using two different guide RNAs (sgATAD3A 7, sgATAD3A 26) compared to non-transduced (NT) cells and cells transduced with one of two empty vectors (EV 3, EV 12). (B) Upregulation of interferon beta (*IFNb*) messenger RNA expression. (C) Upregulation of interferon-stimulated gene (ISG) (*IFI27*, *IFI44L*, *ISG15*, *IFIT1*) messenger RNA expression. Mean values of five independent experiments are shown and are expressed as the ratio of the mRNA levels, normalized to housekeeping gene *HPRT* mRNA, in indicated conditions to that in control EV 3. * indicates significance using a Kruskal Wallis test with Dunn's post-hoc compared to EV 3 for each ISG. (D) Increased phosphorylated IRF3 (pIRF3) (blot on the left, quantification on the right). The specificity of ATAD3A knockdown is indicated by an absence of a change in the expression of either of the mitochondrial proteins TFAM (matrix) or VDAC1 (outer membrane). Quantification of band intensities is expressed as phosphorylated IRF3/total IRF3 ratio and protein levels normalised to Vinculin, averaged across four to five independent experiments, EV is EV 3. (E) Interferon beta (*IFNb*) and representative ISG (*IFI27*, *IFI44L*, *ISG15*, *IFIT1*) mRNA expression upon knockdown of *ATAD3A* with CRISPR editing as in (A) in wild-type (WT) THP-1 cells and in cells null (KO) for either *cGAS*, *STING* or *MAVS*. Mean values of three to five independent experiments are shown and are expressed as the ratio of the mRNA levels, normalized to housekeeping gene *HPRT* mRNA, in indicated conditions to that with EV 3. * indicates statistical significance using 2-way ANOVA with Dunnett's multiple comparison test between cell lines for each CRISPR condition. # indicates statistical significance using 2-way ANOVA with

Dunnett's multiple comparison test between CRISPR conditions for each cell line. (F-G) ISGs, interferon beta (*IFNb*), interferon alpha 2 (*IFNa*) and interferon gamma (*IFNg*) (F), *ATAD3A* and *TFAM* (G) mRNA expression in PBMCs isolated from healthy control donors (HC) or from ATAD3A patients 2 and 4 (AGS1530 and AGS2792 respectively). Mean values and data points of two to three different samples are shown, and are expressed as the ratio of the mRNA levels, normalized to housekeeping gene *HPRT* mRNA, in indicated conditions to that in one HC. * indicates statistical significance in 2-way ANOVA with Dunnett's multiple comparison test to HC for each probe. (H) Representative protein expression analysis of ATAD3A and TFAM expression in PBMCs by western blot. TOM20 is an outer mitochondrial membrane protein, Vinculin a loading control.

**Supplementary Figure S3. Characteristics of interferon signalling induction in ATAD3A patient-derived fibroblasts.**

(A) *ATAD3A* mRNA expression in primary fibroblasts from control human dermal fibroblasts (HDF, average of 3) or from ATAD3A patients 2, 3 and 4 (AGS1530, AGS2216 and AGS2792 respectively). Mean values and data points of ten to fourteen independent experiments are shown, and are expressed as the ratio of the mRNA levels, normalized to housekeeping gene *HPRT* mRNA, in indicated conditions to that in one control HDF. * indicates statistical significance in one-way ANOVA with Dunn's multiple comparison test. (B) Protein expression of ATAD3A in primary fibroblasts assessed by western blot. VDAC1 is a mitochondrial protein, Vinculin a loading control. (C) qPCR analysis of mtDNA copy number per cell, expressed as the ratio of the DNA quantity of the mitochondrial gene (*MT-COXII*) over the nuclear gene (*GAPDH*), in total DNA isolated from fibroblasts. The mean of eight independent experiments is shown. * indicates statistical significance in one-way ANOVA with Dunn's multiple comparison test. (D) Representative ISG (*MX1*, *RSAD2*, *OAS1*, *IFI27*), *cGAS* and *MAVS* mRNA expression, expressed as in (A), in control and ATAD3A patient-derived primary fibroblasts, after downregulation of cGAS (sicGAS) or MAVS (siMAVS) by siRNA or treatment with control siRNA (siCTRL). Mean values and data points of six to nine independent experiments are shown. * indicates statistical significance in 2-way ANOVA with Dunnett's multiple comparison tests between siRNA conditions for each cell line. # indicates statistical significance in 2-way ANOVA with Dunnett's multiple comparison test between control and patient-derived fibroblasts for each siRNA condition. (E) Representative ISG (*Mx1*, *OAS1*, *ISG15*, *IFI27*), *ATAD3A* and *TFAM* mRNA expression, expressed as in (A), in control and ATAD3A patient-derived primary fibroblasts, untreated (UT) or mtDNA-depleted by ddC as in Fig. 4 D. Mean values and data points of three to eight independent experiments are shown. * indicates statistical significance in 2-way ANOVA with Sidak's multiple comparison test between UT and ddC conditions. # indicates statistical significance in 2-way ANOVA with Dunnett's multiple comparison test between control and patient-derived fibroblasts for each treatment. (F) Representative ISG *Mx1* and *ATAD3A* mRNA expression in fibroblasts vehicle-treated (DMSO) or treated with 300 µM 4,4′-diisothiocyanostilbene-2,2′-disulfonic acid (DIDS) or 100 nM rapamycin for 72h. Mean values and data points of five to seven independent experiments are shown. * indicates statistical significance in 2-way ANOVA with Sidak's multiple comparison test between treatment conditions. # indicates statistical significance in 2-way ANOVA with Dunnett's multiple comparison test between control and patient-derived fibroblasts for each treatment. (G) ISG mRNA expression, represented as an ISG score (i.e. the median fold change of mRNA levels of seven ISGs (*RSAD2*, *OAS1*, *Mx1*, *IFI27*, *ISG15*, *IRF7*, *IFI44L*), and BAK1 mRNA expression, expressed as in (A), in fibroblasts, after downregulation of BAX (siBAX) or BAK1 (siBAK1) or both (siBAX+siBAK1) by siRNA or treatment with control siRNA (siCTRL). Mean values and data points of two to three independent experiments are shown. * indicates statistical significance in 2-way ANOVA with Dunnett's multiple comparison tests between siRNA conditions for each cell line. # indicates statistical significance in 2-way ANOVA with Dunnett's multiple comparison test between control and patient-derived fibroblasts for each siRNA condition. (H) Protein BAK1, BAX and ATAD3A levels in primary fibroblasts assessed by western blot, VDAC1 is a mitochondrial protein, Vinculin and Cofilin loading controls. (I) ISG mRNA expression, represented as an ISG score (i.e. the median fold change of mRNA levels of seven ISGs (*RSAD2*, *OAS1*, *Mx1*, *IFI27*, *ISG15*, *IRF7*, *IFI44L*), expressed as in (A), in fibroblasts, after treatment with 50 or 200 µM of BAX inhibitor peptide V5 (BAXi V5) for 72h. Mean values and data points of three to four independent experiments are shown. # indicates statistical significance in 2-way ANOVA with Dunnett's multiple comparison test between control and patient-derived fibroblasts for each treatment.

A Features of systemic sclerosis in patients with mutations in *ATAD3A*.

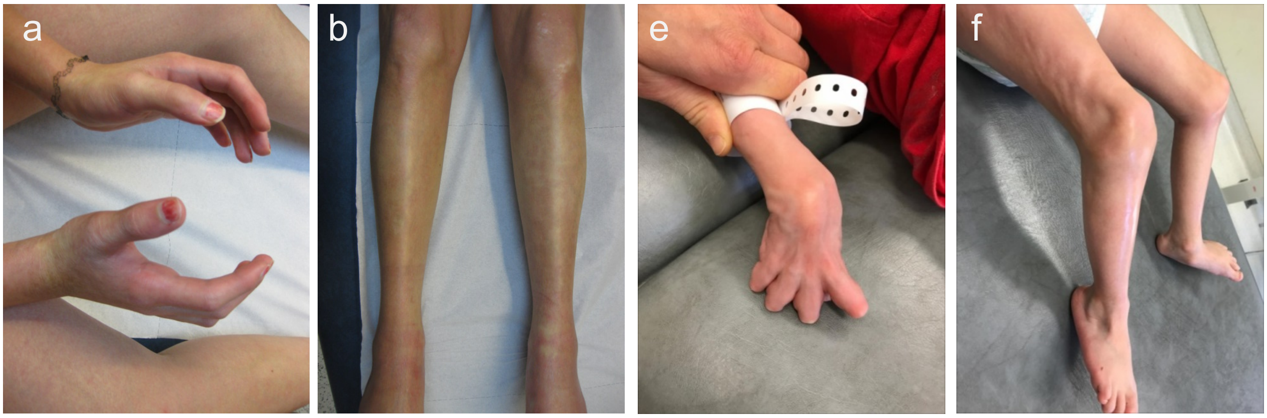


B Histological features of skin biopsies.

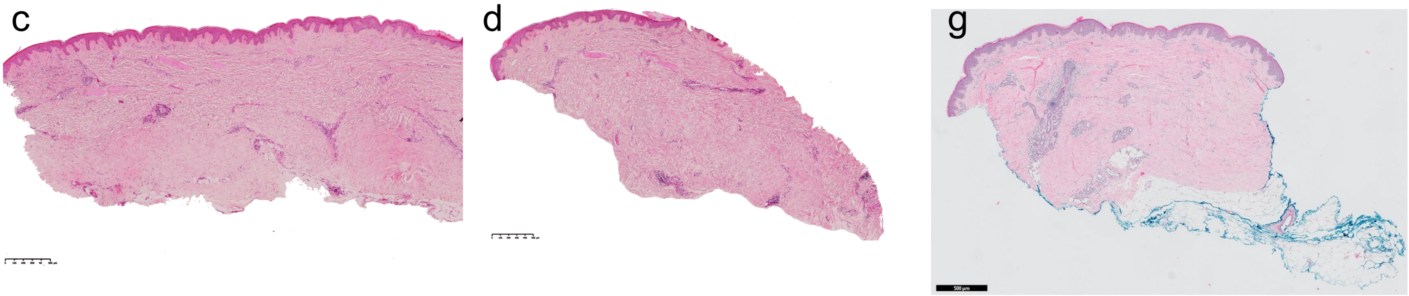


C MRI characteristics.

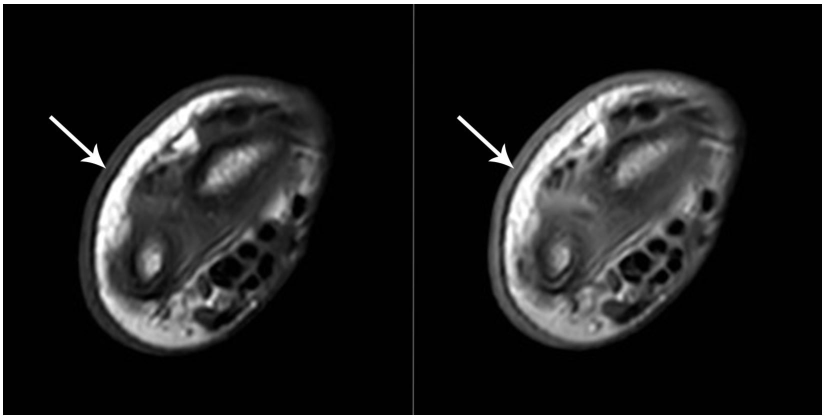

**Supplementary Figure S1.** *Lepelley et al.*

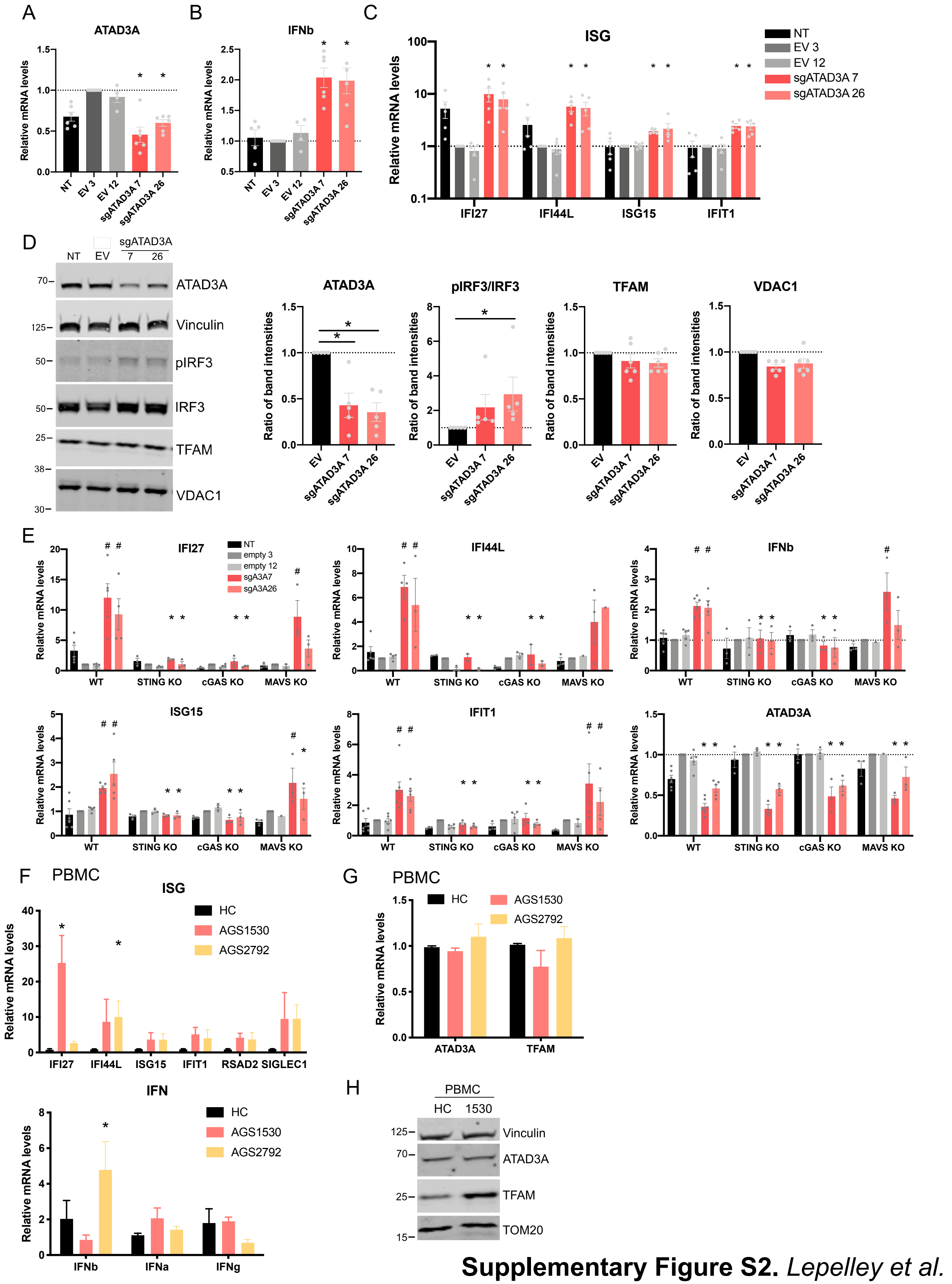


**Supplementary Figure S2.** *Lepelley et al.*

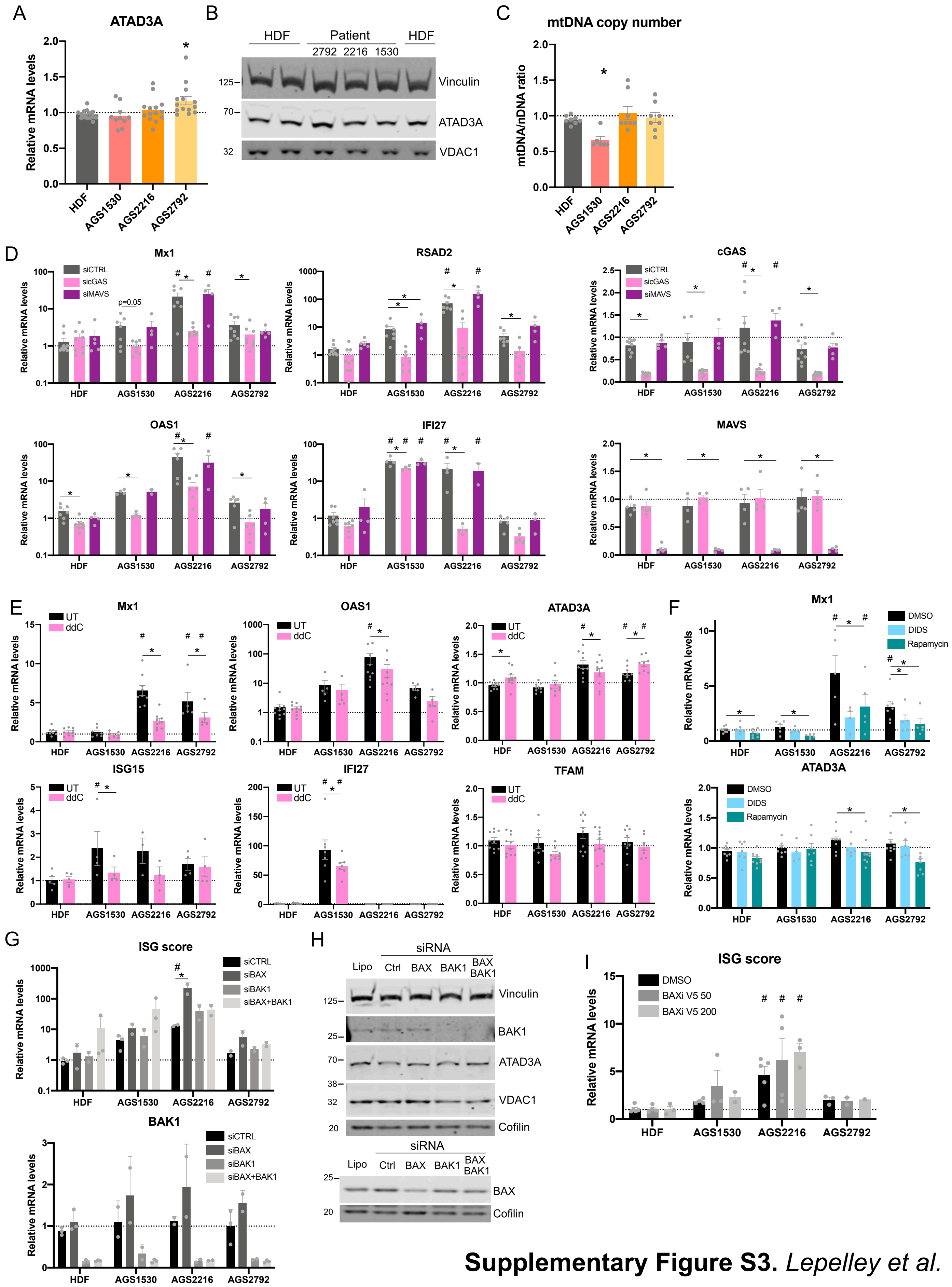


**Supplementary Figure S3.** *Lepelley et al.*

**Supplementary Table 1**. Features related to systemic sclerosis in Patient 2 and Patient 3.

| | Patient 2 | Patient 3 |
|---|---|---|
| **Age at presentation** | 21 years | 2 years |
| **History** | Pain and swelling of fingers, with the rapid development of scleroderma of the hands, face and ventral surface of the forearms | Reddish-blue indurated skin lesions initially involving the lower limbs. Progressive hardening of skin and change in facial appearance due to a loss of fat tissue. Myalgia and arthralgia causing unrest, particularly at night. At age 4 years: perioral sclerosis with open mouth and rigidity of the temporomandibular joints, and tight, hard, sclerodermatous skin of the upper and lower limbs, less marked over the abdomen |
| **Contractures** | Knees, ankles, wrists and fingers | Knees, ankles, elbows, wrists and fingers |
| **Rodnan skin score** | 33/51 | 32/51 |
| **Skin biopsy** | Massive collagen fibrosis with loss of elastic fibers and a lymphocytic infiltrate. No immunoglobulin or complement deposition | Mild lymphocytic infiltration in the superficial dermis, with deep dermal sclerosis and loss of skin appendages surrounded by thick collagen bundles |
| **Capillaroscopy** | Normal | Normal |
| **Antinuclear antibodies (titer)** | Persistently elevated (1/5220) | Absent |
| **ENA antibodies (titer)** | Persistently elevated (1/5220) | Absent |
| **Anti ds-DNA antibodies (titer)** | Persistently elevated (1/800) | Absent |
| **Anti-SCL70; anti-centromere; anti-fibrillarin; anti-RNA polymerase III antibodies** | Absent | Absent |
| **Gamma-globulins (normal < 13g/l)** | Persistently elevated (25g/l) | Normal |
| **C-reactive protein (normal < 6mg/l)** | Minimally elevated (11mg/l) | Not elevated |
| **Limb magnetic resonance imaging** | Not performed | Cutaneous thickening in the upper limbs, and mild contrast enhancement of subcutaneous fat tissue without calcifications |
| **Ulcerations** | No | No |

| | | |
|---|---|---|
| **Calcinosis** | No | No |
| **Raynaud's** | No | No |
| **Fever** | No | No |
| **Response to therapy** | Steroids associated with remission of arthralgia. Rapamycin associated with partial regression of cutaneous fibrosis (Rodnan score: 10/51) | No response to non-steroidals. Methylprednisolone and rapamycin associated with resolution of nocturnal unrest and cessation of pain medication, with some clinical gains including reduced induration of sclerotic lesions and improved mouth closure and feeding |
| **Current therapy** | Rapamycin | Rapamycin and low-dose methylprednisolone |
